# Persistence of a novel regeneration-associated transitional cell state in pulmonary fibrosis

**DOI:** 10.1101/855155

**Authors:** Yoshihiko Kobayashi, Aleksandra Tata, Arvind Konkimalla, Hiroaki Katsura, Rebecca F. Lee, Jianhong Ou, Nicholas E. Banovich, Jonathan A. Kropski, Purushothama Rao Tata

**Affiliations:** Department of Cell Biology, Duke University School of Medicine, Durham, NC 27710, USA; Medical Scientist Training Program, Duke University School of Medicine, Durham, NC 27710, USA; Translational Genomics Research Institute, Phoenix, AZ 85004, USA; Division of Allergy, Pulmonary and Critical Care Medicine, Department of Medicine, Vanderbilt University Medical Center, Nashville, TN 37212, USA; Department of Veterans Affairs Medical Center, Nashville, TN 37212, USA; Department of Cell and Developmental Biology, Vanderbilt University, Nashville, TN 37212, USA; Duke Cancer Institute, Duke University School of Medicine, Durham, NC 27710, USA; Regeneration Next, Duke University, Durham, NC 27710, USA

## Abstract

Stem cell senescence is often seen as an age associated pathological state in which cells acquire an abnormal and irreversible state. Here, we show that alveolar stem cell differentiation during lung regeneration involves a unique previously uncharacterized transitional state that exhibits cardinal features normally associated with cell senescence. Specifically, using organoid cultures, multiple *in vivo* injury models coupled with single cell transcriptomics and lineage tracing analysis, we find that alveolar stem cell differentiation involves a novel, pre-alveolar type-1 transitional state (PATS) en route to their terminal maturation. PATS can be distinguished based on their unique transcriptional signatures, including enrichment for TP53, TGFβ, and DNA damage repair signaling, and cellular senescence in both *in vivo* and *ex vivo* regenerating tissues. Significantly, PATS undergo extensive cell stretching, which makes them vulnerable to DNA damage, a feature commonly associated with most degenerative lung diseases. Importantly, we find enrichment of PATS-like state in human fibrotic lung tissues, suggesting that persistence of such transitional states underlies the pathogenesis of pulmonary fibrosis. Our study thus redefines senescence as a state that can occur as part of a normal tissue maintenance program, and can be derailed in human disease, notably fibrosis.

## Introduction

Adult stem cells undergo dynamic changes in phenotype in response to tissue damage ^1, 2^. These changes include resurgence from a quiescent or poised state, onset of proliferation, activation of new programs of gene expression and a return to homeostasis ^3–8^. In many cases, repair also involves changes in epithelial cell shape, for example through transient stretching and expansion to cover areas of damage or denudation, or more permanent changes in cell phenotype. Studies of stem cells during regeneration usually focus on understanding how cells select new differentiation programs in response to signals from the niche. However, much less is known about the significance of changes in parameters such as cell shape and spreading and whether they involve the transient and tightly controlled expression of genes usually associated with pathologic states such as DNA repair and senescence. Here, we explore this question in relation to epithelial repair in the distal gas-exchange alveolar region of the mammalian lung.

In the lung, the maintenance of the alveolar epithelium at homeostasis and its regeneration after injury are fueled by surfactant-producing, cuboidal, type-2 alveolar epithelial cell (AEC2), which can self-renew and differentiate into very large, thin, type-1 alveolar epithelial cells (AEC1), specialized for gas exchange ^1, 2, 9–17^. Recent studies have identified a subset of AEC2s that are enriched for active Wnt signaling and have higher “stemness” compared to Wnt-inactive AEC2s. Such differences in alveolar progenitor cell subsets have been attributed to differences in microenvironmental signals ^9, 12, 18^; in this case, to the vicinity of PDGFRα-expressing alveolar fibroblasts which produce ligands to activate Wnt signaling in AEC2s. Recent studies have also implicated other intercellular signaling pathways, including BMP, Notch, TGFβ, YAP, and NF-κB, in the proliferation and differentiation of AEC2s, both at steady state and in response to alveolar injury ^15, 19–21^. However, the precise mechanisms by which the cuboidal AEC2s orchestrate their dramatic changes in cell shape, structure and mechanical properties as they convert into thin, flat AEC1s, remain elusive. In addition, the cellular mechanisms that drive AEC2s to express genes associated with cell senescence, a feature commonly observed in most progressive pulmonary diseases, remain unknown.

Here, using organoid cultures and single-cell transcriptome studies, we uncovered novel, distinct cell states encompassing the transition between AEC2s and AEC1. Moreover, murine lineage tracing, coupled with injury repair models, have revealed the existence of similar transition states *in vivo*. Our study reveals signaling pathways that control these transition states. We also discovered that these novel transitional states exhibit DNA damage responses, and express senescence-related genes en route to AEC1. Importantly, these transitional states correlate with abnormal epithelial cells that are associated with defective fibrotic foci in lungs of human patients with progressive pulmonary fibrosis.

## Results

### Single-cell transcriptomics revealed novel alveolar epithelial cell states in *ex vivo* organoids

Recent studies have shown that in response to lung injury, AEC2s proliferate and give rise to AEC1s ^11^. Moreover, in alveolar organoid culture AEC2s spontaneously generate AEC1s ^10^. However, the molecular mechanisms and transitional cell states underlying the differentiation of AEC2 into AEC1 is still poorly understood. To understand the mechanisms associated with AEC2 differentiation to AEC1, we performed single-cell transcriptome analysis on cells isolated from alveolar organoids. Purified AEC2 were mixed with fibroblasts and cultured for 10 days to grow organoids for scRNA-seq analysis (Fig. 1a). Uniform Manifold Approximation and Projection (UMAP) identified two major clusters consisting of *Epcam*^+^ epithelial cells and *Vim*^+^/*Pdgfra^+^* fibroblasts (Supplementary Fig. 1a,b). Next, we further deconvoluted and visualized epithelial cell populations. Within the epithelial cells, we observed multiple sub-clusters: cells expressing high levels of *Sftpc,* a marker for AEC2s, cells expressing high levels of *Ager* (*Rage*), a marker for AEC1s, and *Sftpc^+^/Mki67^+^* proliferating AEC2s (Fig. 1b). These data indicate that under these culture conditions, purified lineage labeled AEC2s proliferate and spontaneously generate AEC1s. In addition, we identified a novel population of alveolar epithelial cells expressing *Cldn4*, *Krt19*, and *Sfn* (Fig. 1b,c). The marker genes unique to this cluster showed two distinct patterns when visualized in UMAP and volcano plots (Fig.1 c,d). One subset (*Ctgf^+^* cells) is enriched for *Ctgf, Clu, Sox4 and Actn1* while the other (*Lgals3*^+^ cells) is enriched for *Lgals3, Csro1, S100a14, and Cldn18* (Fig.1 c,d). Additional transcripts that are enriched in the *Lgals3^+^* sub-cluster include *Ager, Emp2,* and *Hopx,* markers of AEC1, suggesting resemblances and a potential lineage hierarchy between *Lgals3^+^* cells and AEC1 (Fig. 1c,d). These data suggest that the newly identified *Cldn4/Krt19/Sfn*^+^ population is an intermediate between AEC2 and AEC1. We therefore termed this population “<u>p</u>re-<u>a</u>lveolar type-1 transitional cell <u>s</u>tate” (in short, PATS). To validate our single cell data, we performed immunofluorescence analysis for PATS markers on alveolar organoids (Fig. 1e). Immunostaining analysis confirmed the presence of cells expressing PATS-specific markers, including CLDN4, LGALS3, and SOX4 in alveolospheres (Fig. 1f). Taken together, these data identified unique and novel cell states during alveolar epithelial stem cell differentiation in organoid cultures.

**Figure 1.**
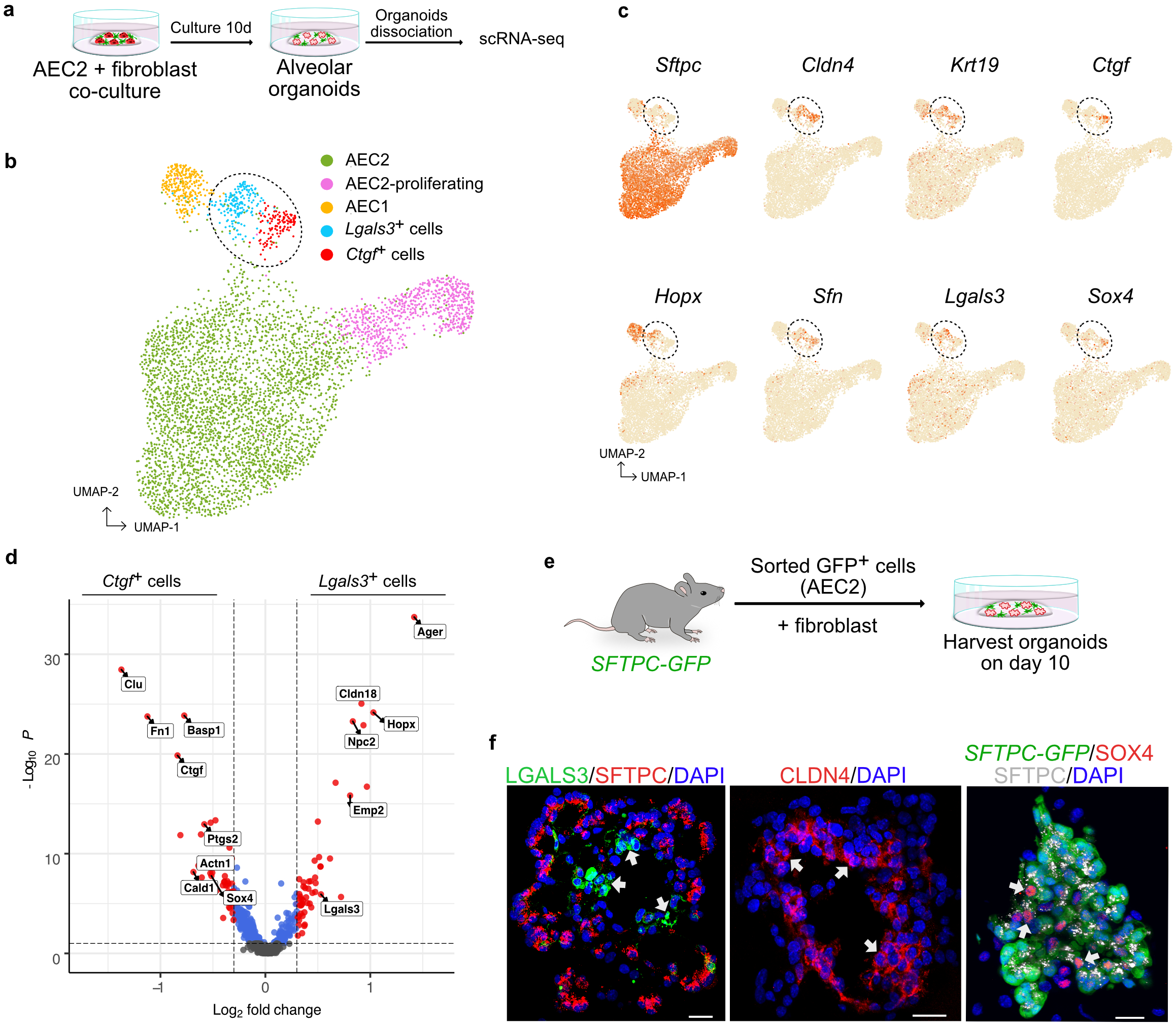
scRNA-seq reveals novel alveolar epithelial cell states in *ex vivo* organoids. a, Schematic of alveolar organoid culture utilized for single-cell RNA-seq. b, Uniform manifold approximation and projection (UMAP) visualization of epithelial populations in cultured alveolar organoids. AEC2 (green) – alveolar epithelial type-2 cells, AEC2-proliferating (pink) – proliferating alveolar epithelial type-2 cells, AEC1 (yellow) – alveolar epithelial type-1 cells, New cell states :*Lgals3*^+^ cells (blue) and *Ctgf*^+^ cells (red) c, UMAP plots show the expression of indicated genes in epithelial populations in cultured alveolar organoids. Dotted circles in b and c indicate the novel cell states. d, Volcano plot shows specific genes to *Ctgf*^+^ population and *Lgals3*^+^ population. e, Schematic of alveolar organoid culture using fibroblasts and AEC2 cells sorted from *SFTPC-GFP* mice. f, Immunostaining for PATS markers in alveolar organoids. Co-staining of LGALS3 (green) and SFTPC (red) is shown in left panel; localization of CLDN4 (red) - middle panel and *SFTPC*-GFP (green) co-stained with SOX4 (red) and SFTPC (grey) (right panel). DAPI stains nuclei (blue). Scale bars indicate 20 μm. Arrows indicate PATS.

### PATS emerge *in vivo* after alveolar injury

We then asked whether PATS can be observed *in vivo* in homeostatic and regenerating alveolar tissues. To test this, first we analyzed a publicly available scRNA-seq data set from alveolar epithelial cells isolated from mice exposed to lipopolysaccharide (LPS), a bacterial endotoxin that causes lung injury ^22^. We rendered scRNA-seq data from LPS and control lungs in UMAP plots and found a population that is unique to LPS treated lungs but is not in controls. Significantly, this population is enriched for genes expressed in PATS, including *Cldn4, Sox4, Lgals3,* and *Fn1* (Supplementary Fig. 2). To further characterize PATS, we utilized *Ctgf-GFP* mice, a fluorescence reporter transgenic mouse line in which green fluorescent protein is driven by the promoter for *Ctgf*, a gene highly enriched in PATS ^23^. We then exposed these mice to bleomycin, a drug that damages the alveolar region causing transient fibrosis. We collected lungs on day 12 post injury for analysis (Fig. 2a). Immunofluorescence analysis for GFP expression in control, uninjured, *Ctgf-GFP* mice revealed GFP signal specifically in alveolar fibroblasts and not in alveolar epithelial cells (Fig. 2b,c and Supplementary Fig. 3). By contrast, in bleomycin injured lungs, we found GFP expression in a significant number of epithelial cells co-labeled by the expression of the AEC2 marker, SFTPC (Fig. 2b,c and Supplementary Fig. 3). These data corroborate our organoid scRNA-seq data and suggest that PATS arise from AEC2s after alveolar injury. Next, we performed co-immunofluorescence analysis for other PATS markers including CLDN4, LGALS3 and SFN. Our analysis revealed that *Ctgf*-GFP expression in epithelial cells overlaps with that of other PATS markers in bleomycin-injured mice but not in control lungs (Fig. 2 b,c). Interestingly, we noticed that many cells that are positive for *Ctgf*-GFP and other PATS markers show an elongated cell shape as compared to cuboidal AEC2, suggesting that they are en route to differentiation into AEC1 (Fig. 2b,c). Flow cytometric analysis further revealed a significant number of *Ctgf*-GFP^+^ cells and LGALS3^+^ cells in bleomycin treated lungs compared to controls (Fig. 2d). A recent study found elevated levels of KRT8 expression in alveolar epithelial cells after bleomycin-induced injury ^24^. Therefore, we tested the expression of KRT8 in alveoli after bleomycin-induced lung injury. We found a significant increase in overlap between KRT8 and *Ctgf*-GFP-expressing cells (Supplementary Fig. 3). Taken together, our data reveal that cells with a PATS phenotype emerge in alveoli after bleomycin injury *in vivo*.

**Figure 2.**
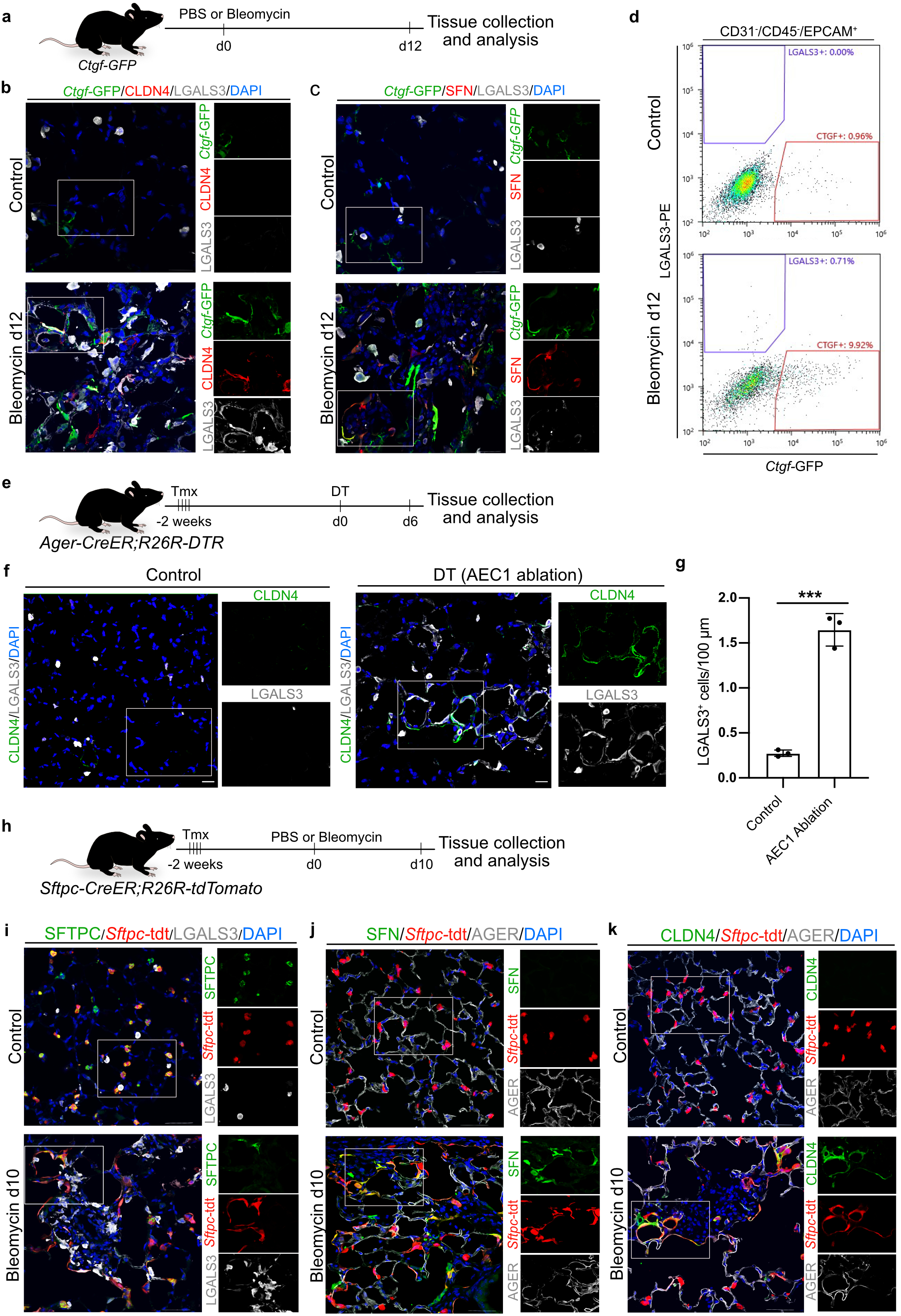
The novel alveolar cell states emerge transiently and originate from alveolar stem cells after alveolar injury *in vivo*. a, Schematic of bleomycin-induced lung injury in *Ctgf-GFP* mice. b-c, Immunostaining for PATS markers. b, co-staining for *Ctgf*-GFP (green), CLDN4 (red) and LGALS3 (grey), c, Co-staining for *Ctgf*-GFP (green), SFN (red) and LGALS3 (grey) in control lung (upper) and bleomycin-treated lungs collected on day 12 after injury (lower). d, Flowcytometric analysis of PATS in *Ctgf*-GFP mouse model. The percentage of *Ctgf*-GFP (CD31^-^/CD45^-^ /EPCAM^+^) (red line) and LGALS3 (CD31^-^/CD45^-^/EPCAM^+^) (blue line) are shown as indicated in control (upper panel) and bleomycin injures (lower panel). e, Experimental design to ablate AEC1 cells in *Ager-CreER;R26R-DTR* mouse model. Mice were administered with tamoxifen (Tmx) followed by diphtheria toxin (DT) and tissue collection on day 6. f, Immunostaining for CLDN4 (green) and LGALS3 (grey) in control (left) and AEC1-ablated lungs (right). g, Mean linear intercept length analysis of LGALS3^+^ cells in control and AEC1-ablated lungs. Asterisks indicate *p* < 0.0002 (un-paired student’s t-test). Data are from three independent experiments and are presented as mean ± s.e.m. h, Experimental workflow for sequential administration of tamoxifen followed by bleomycin injury and tissue collection for analysis using *Sftpc-CreER;R26R-tdTomato* mice. i-k, Immunostaining for PATS markers in Sftpc-lineage labeled cell in control (upper panel) and bleomycin inured lungs (lower panel). i, SFTPC (green), *Sftpc*-tdt (red) and LGALS3 (grey), j, SFN (green), *Sftpc*-tdt (red) and AGER (grey), and k, CLDN4 (green), *Sftpc*-tdt (red) and AGER (grey). DAPI stains nuclei (blue). Scale bars indicate 30 µm. White line box in merge image indicates region of single channel images shown in left side.

We then asked whether PATS cells are specific to bleomycin injury or appear in other injury models. To test this we employed *AGER-CreER;R26R-DTR* mice in which administration of tamoxifen activates the expression of diphtheria toxin receptor in AGER-Expressing AEC1 cells. This receptor then binds to exogenously administered diphtheria toxin resulting in selective ablation of AEC1 cells (Fig. 2a). We collected lung samples at day 6 after DT injection and performed immunostaining analysis for PATS markers (Fig. 2e). Interestingly, we found cells expressing CLDN4 and LGALS3 that appear to show elongated morphology in AEC1 ablated lungs but not in controls (Fig. 2f and Supplementary Fig. 4). Further quantification revealed a significant number of LGALS3^+^ cells after AEC1 ablation (1.647±0.1041 after ablation vs 0.274± 0.02086 in control) (Fig. 2g). These data suggest that emergence of PATS can be a general mechanism in alveolar regeneration.

### Lineage tracing revealed PATS cells originate from AEC2

Our above data from *ex vivo* organoid cultures and *in vivo* injury models suggested that PATS cells originate from AEC2s (Fig. 1). To empirically test this hypothesis, we used a *Sftpc-creER;R26R-tdTomato* mouse model, in which tamoxifen administration permanently induces the expression of tdTomato fluorescent protein specifically in AEC2 and their descendants (Fig. 2h) ^25^. On day-10 post bleomycin administration, we observed lineage labeled (tdTomato^+^) cells co-expressing LGALS3 (Fig. 2i), SFN (Fig. 2j) and CLDN4 (Fig. 2k), and high levels of KRT8 (Supplementary Fig. 5) in damaged regions of the alveoli. Of note, LGALS3, SFN and CLDN4 expression was not observed in tdTomato+ cells in the control lungs (Fig. 2i-k). Interestingly, some lineage labeled SFN^+^ and CLDN4^+^ cells showed flat, thin, and elongated morphology and co-expressed AGER, implying that they are in transition to AEC1. Taken together, our data provide evidence that PATS cells emerge *in vivo* from AEC2s after alveolar injury and generate AEC1 (Fig. 2).

### Lineage tracing analysis revealed PATS cells generate AEC1

Multiple lines of data from our above experiments suggest that PATS are en route to AEC1 (Fig. 1 and Fig. 2). To test this possibility, we first used an algorithm tool called Velocyto^26^, which allows the prediction of cell differentiation trajectories based on ratios between spliced and unspliced mRNA. Using this algorithm, we observed that strong RNA velocities (trajectory) originating from the newly identified *Ctgf*^+^ population to AEC1 through the *Lgals3*^+^ population (Fig. 3a-c). These data further strengthened our above findings that PATS cells are intermediate between AEC2 and AEC1. We then used *Krt19-CreEr an allele of* a gene specifically expressed in PATS, in combination wit*h R26R-tdTomato* (in short, *Krt19-tdTomato*) to carry out lineage tracing in the bleomycin lung injury model to test whether the same trajectory occurs in regenerating alveoli *in vivo*. First, mice were exposed to bleomycin to induce alveolar injury, then tamoxifen was subsequently administered on day 7 after the exposure to label *Krt19*-expressing cells and their progeny by tdTomato (Fig. 3d). In control lungs, we observed a small number of tdTomato labeled cells co-expressing AGER^+^ (1.39%±0.31%) and LGALS3^hi^ macrophages (Fig. 3g and Supplementary Fig. 6a,c). However, we did not find any labeling in AEC2 cells (0%±0) (Supplementary Fig. 6a,c). In contrast, we observed a significant number of tdTomato^+^ cells co-expressing low levels of SFTPC in damaged regions in bleomycin injured lungs (47.22%±6.81%) (Supplementary Fig. 6 a-c). Of note, we did not find tdTomato^+^ cells co-expressing SFTPC in uninjured regions from the same lungs, indicating that *Krt19* expression is specifically activated in response to injury in the damaged regions. These data also indicate that the *Krt19-tdTomato* mouse line and the conditions we tested here are well suited for labeling and tracing PATS. Furthermore, co-immunofluorescence analysis revealed lineage labeled cells co-expressed PATS markers in bleomycin treated lungs but not in control lungs (Fig. 3e-g and Supplementary Fig. 6). Specifically, we observed SFN expression in tdTomato^+^ cells that appear cuboidal (white arrows) as well as in elongated cells (yellow arrowheads) (Fig. 3e-g, right). By contrast, CLDN4 and LGALS3 expression was present mostly in elongating cells (yellow arrowhead and white arrow, respectively)) (Fig. 3e-g right). Importantly, on day-12 post bleomycin administration, we found numerous *Krt19*-tdTomato^+^ cells co-expressing AGER, a marker for AEC1 (Fig. 3e,f and Supplementary Fig. 6). Thus, our combined RNA velocity and lineage tracing analysis reveal that the newly identified PATS traverse between AEC2 and AEC1.

**Figure 3.**
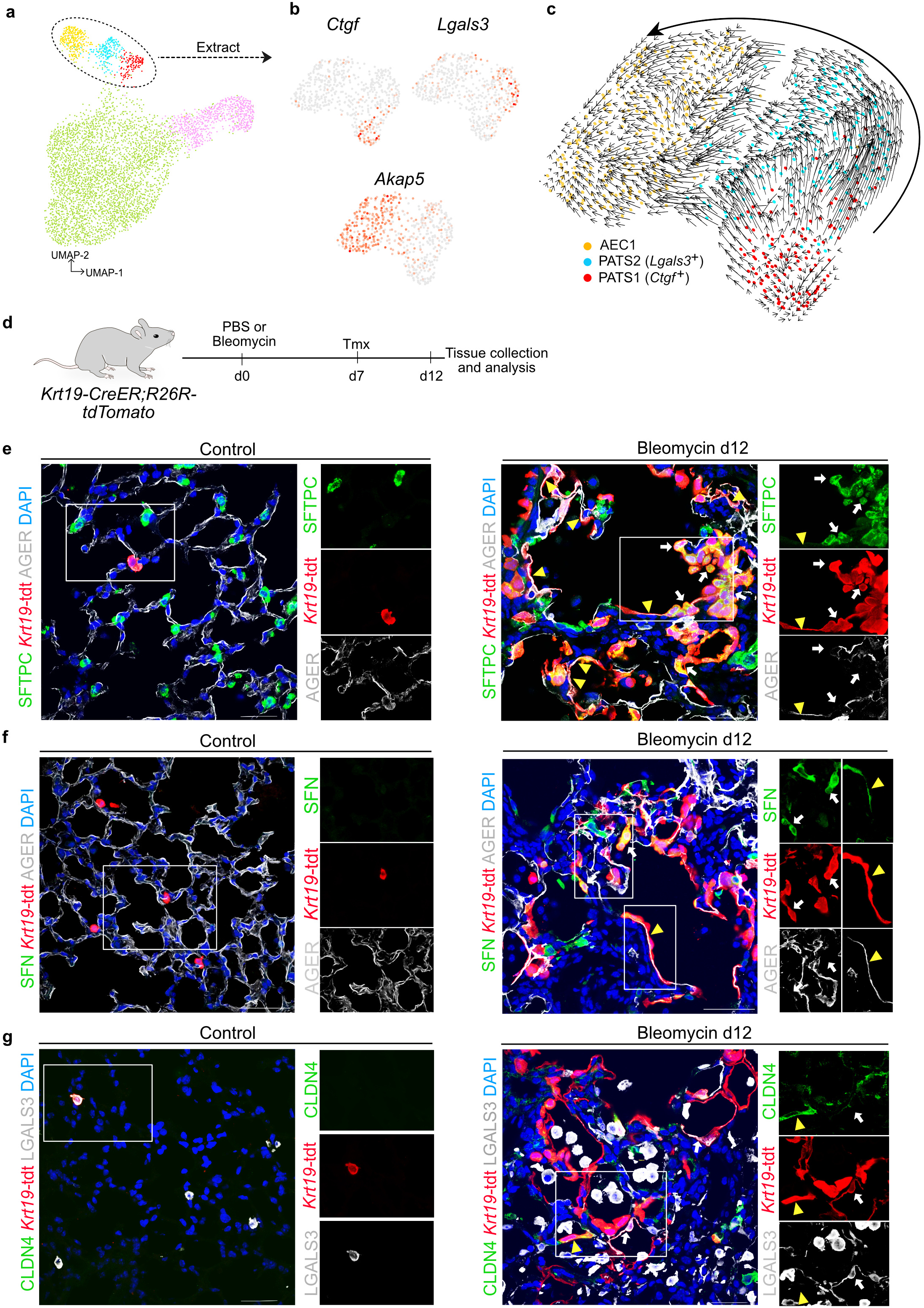
Lineage tracing revealed that PATS generate AEC1. a, UMAP of epithelial populations in cultured alveolar organoids. Arrow indicates selected cell populations in the oval-shaped circle are shown in panel b. b, UMAP plots show the expression of indicated genes in the selected populations (oval-shaped circle in panel a). c, RNA velocity analysis for PATS and AEC1. Arrows indicate predicted lineage trajectories. d, Schematic representation of experimental design to sequentially administer bleomycin (injury) or PBS (control) followed by tamoxifen (to label Krt19-expressing cells) in *Krt19-CreER;R26R-tdTomato* mice. e. Immunostaining for SFTPC (green), *Krt19*-tdt (red) and AGER (grey). White arrows indicate SFTPC^+^, *Krt19*-tdt^+^ cells. Yellow arrowhead indicates AGER^+^, *Krt19*-tdt^+^ cells. (Scale bar: 30 µm). f, Co-staining for SFN (green), *Krt19*-tdt (red) and AGER (grey). White arrows indicate SFN^+^, *Krt19*-tdt^+^ cells. Yellow arrowhead indicates SFN^+^, *Krt19*-tdt^+^, AGER^+^ cell. (Scale bar: 50 µm). g. Immunostaining for CLDN4 (green), *Krt19*-tdt (red) and LGALS3 (grey). White arrows indicate CLDN4^+^, *Krt19*-tdt^+^ cells. Yellow arrowhead demonstrates LGALS3^+^, *Krt19*-tdt^+^ cell. (Scale bar: 30 µm). e-g Control lungs are shown in left panels and bleomycin day 12 injured lungs are shown in right panels. DAPI stains nuclei (blue). White line box in merge image indicates region of single channel images shown in left side.

### Conserved Transcriptional programs and pathways in PATS

Differentiation of AEC2 to AEC1 is associated with a dramatic change in the shape and structure of the cells, from a cuboidal to flat and thin morphology. Such a transition is typically accompanied by many changes in the expression of signaling and structural proteins ^27, 28^. We therefore hypothesized that these changes occur in PATS. To test our hypothesis, we analyzed our scRNA-seq data from *ex vivo* organoid cultures and *in vivo* alveolar injury models (LPS-induced lung injury), focusing on genes that were commonly enriched in both datasets. We found numerous genes that are conserved among PATS (Fig. 4a). Pathway enrichment analysis revealed activation of TP53, TNF/NF-κB, ErbB, HIF1, cell cycle arrest, cytoskeletal dynamics, tight junction signaling and TGFβ signaling in PATS cells compared to other populations (Fig. 4 b,c). Previous studies have implicated all these pathways in alveolar regeneration following injury, highlighting the importance of these pathways in PATS ^20, 22, 29–31^. Unexpectedly, we also found significant enrichment for genes representative of senescence and DNA damage response pathways (Fig. 4).

**Figure 4.**
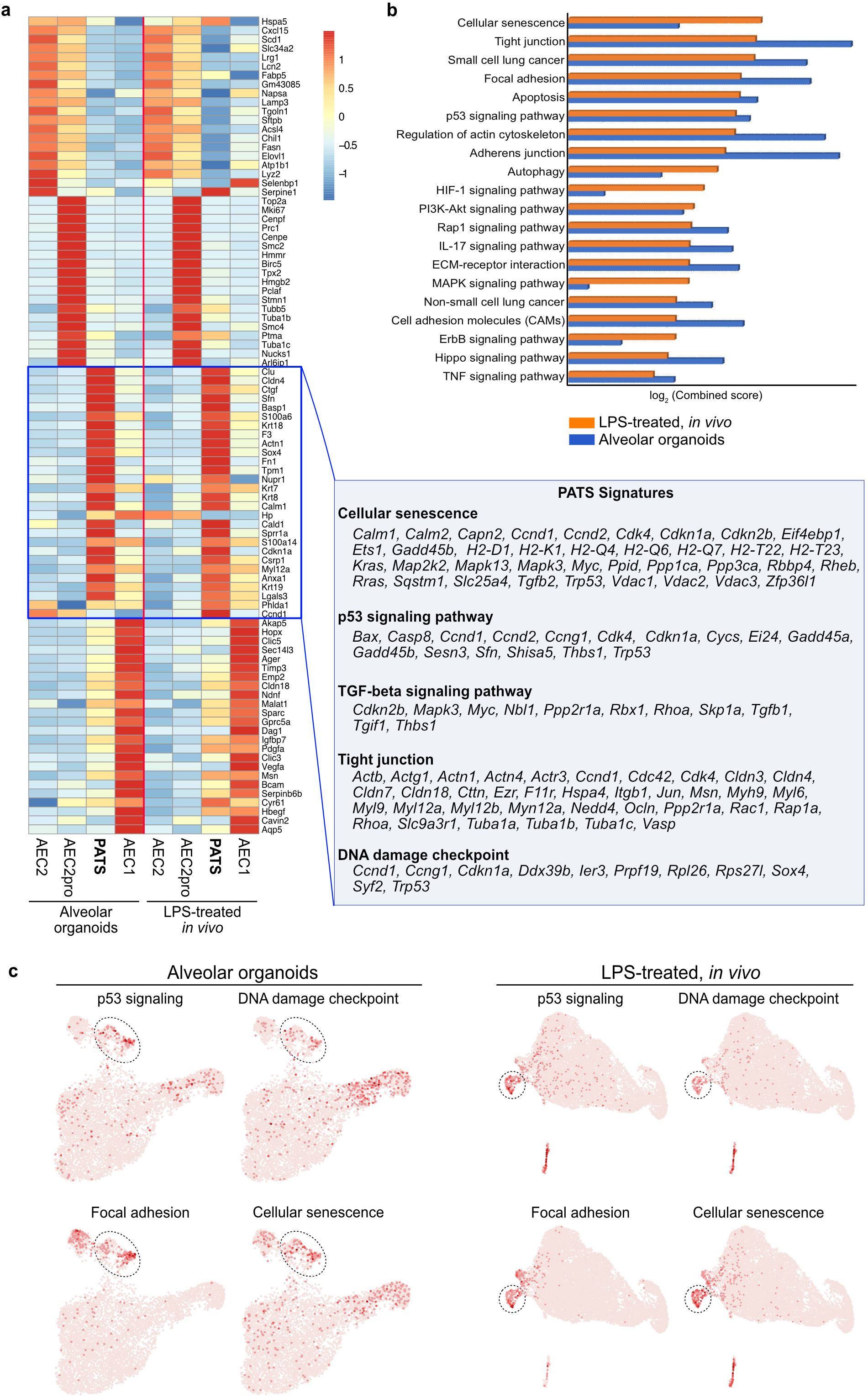
Gene expression signatures and signaling pathways enriched in PATS. a, Heatmap shows marker gene expression of each cell population in alveolar organoids (left) and in LPS-treated murine lungs (right) (scale shows z-score). Table on the right indicates genes enriched in the indicated pathways and cellular processes specifically in PATS. b, KEGG pathways enriched in PATS. Scale shows log_2_ (combined score) obtained from web-based tool - Enrichr. c, UMAP rendering of PATS enriched signaling pathways and cellular processes in alveolar organoids (left) and LPS-treated murine lungs (right).

### PATS cells naturally exhibit DNA damage and senescence during alveolar regeneration *in vivo*

Our above data indicated that DNA damage and senescence associated genes are highly enriched in PATS. To validate these findings, we assayed lung tissue sections from bleomycin or PBS treated mice for *β-galactosidase* activity, that serves as a biomarker for senescent cells, as well as γH2AX, a marker for DNA damage signaling. Interestingly, we detected numerous *β-galactosidase* active cells, as assessed based on X-gal derived blue color deposition, in *Sftpc*-tdt lineage derived cells in bleomycin injured alveoli but not in control alveoli (Fig. 5a,b). Similarly, we also observed accumulation of γH2AX puncta in the nuclei of elongated *Sftpc*-tdt^+^ cells (Fig. 5c,d). Moreover, quantification revealed that 15.09%±1.52% *Sftpc*-tdt^+^ cells expressed γH2AX in bleomycin injured lungs compared to 0.84%±0.25% in controls (Fig. 5d). Furthermore, we found that γH2AX^+^ cells also co-expressed LGALS3, indicating that PATS undergo DNA damage during alveolar regeneration.

**Figure 5.**
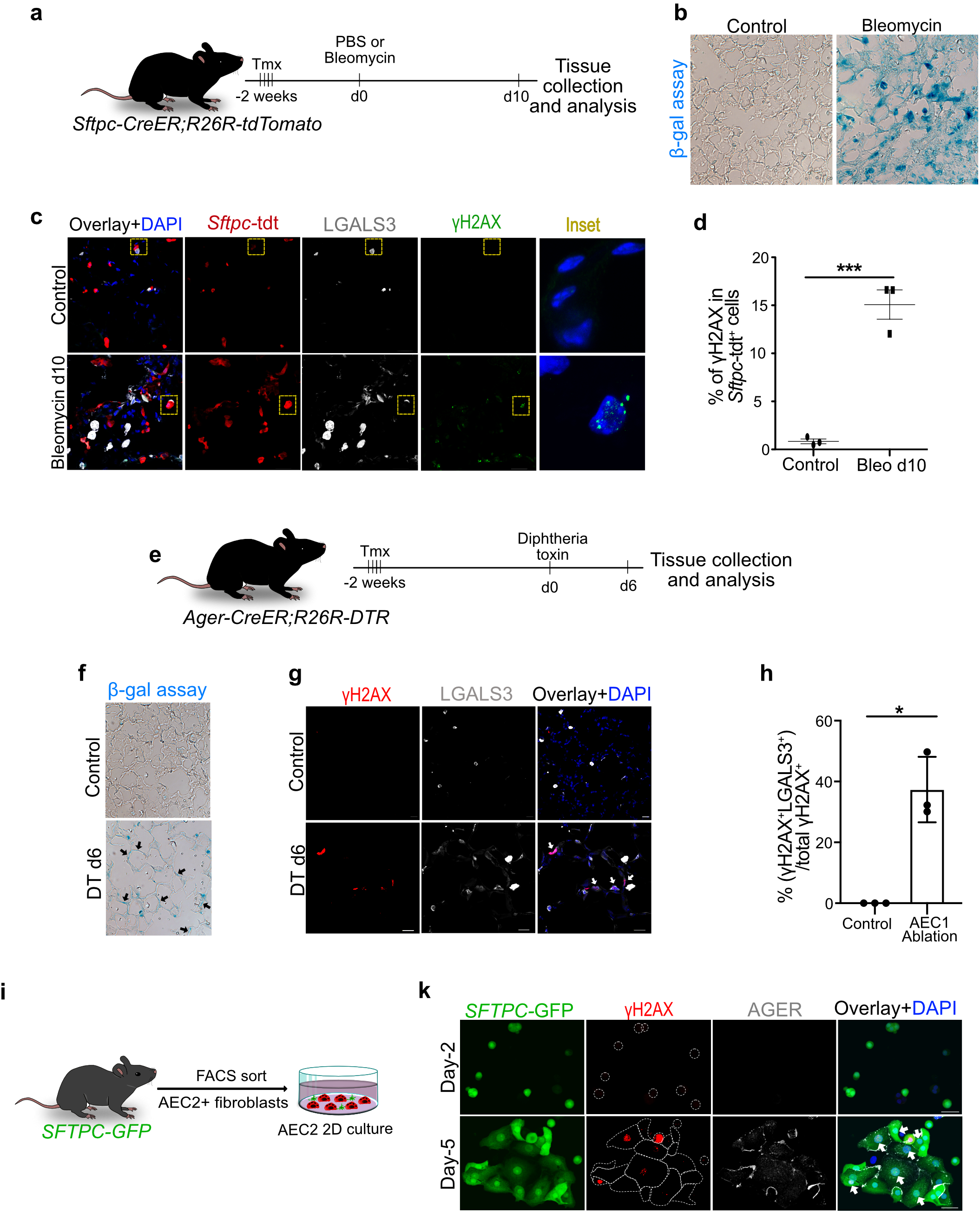
PATS undergo stretch-induced DNA damage *in vivo* and *ex vivo*. a, Schematic of bleomycin-induced lung injury in *Sftpc-CreER;R26RtdTomato* mice. Immunostaining for b, β-galactosidase staining in bleomycin injured and control lungs. c, γH2AX (green), *Sftpc*-tdt (red) and LGALS3 (grey) in control (upper) and bleomycin-treated lung collected on 12 days after the injury (lower). Inset on the right side in c shows the higher magnification of region indicated by dotted yellow box. DAPI stains nuclei (blue). Scale bar: 30 µm. d, Quantification of γH2AX^+^ cells in *Sftpc*-tdt^+^ cells in control and bleomycin day 10 injured mice. Asterisks indicate *p* < 0.0008 (un-paired student’s t-test). Data are from three independent experiments and are presented as mean ± s.e.m. e, Schematic of AEC1 ablation using diphtheria toxin (DT). f, β-galactosidase staining in DT-treated and control lungs. g, Co-staining for γH2AX (red) and LGALS3 (grey) in control (left) and DT-treated lung collected on 6 days after the injury (right). White arrows indicate γH2AX^+^, LGALS3^+^ cells. Scale bar 20 µm h, Quantification of LGALS^+^/γH2AX^+^ cells in total γH2AX^+^ cells in control and DT-treated lung. Asterisk indicates *p* = 0.05. Data are from three independent experiments and are presented as mean ± s.e.m. i, Schematic of alveolar organoid culture. j, Co-staining for *SFTPC*-GFP (green) and γH2AX (red) in alveolar organoids. Inset on the right side shows the higher magnification of region indicated by dotted yellow box. White arrowheads indicate γH2AX^+^ cells Scale bar: 30 µm. k, Schematic of 2D culture of AEC2. l, Immunostaining for *SFTPC*-GFP (green), γH2AX (red) and AGER (grey) in 2D culture of AEC2.

Bleomycin is known to induce DNA damage in cells. However, both our scRNA-seq analysis and immunostaining revealed a strong enrichment for DNA damage repair signaling in PATS but not in other cell types. To avoid any potential effects from bleomycin on DNA damage, we performed γH2AX staining on lungs from the AEC1-specific cell ablation model. In this model, 37.37%±6.197 LGALS3^+^ cells showed γH2AX^+^ nuclear speckles in AEC1 cells compared to 0%±0% in control lungs (Fig. 5e-h). Of note, γH2AX^+^ cells were not observed in experimental lungs after recovery of AEC1 cells, suggesting that the DNA damage observed in PATS cells is transient and repaired as the cells progress to AEC1 (data not shown).

### PATS cells are vulnerable to mechanical stretch-induced DNA damage

Previous studies have shown that ionizing radiation, oxidative stress, and the mechanical stretching that occurs during cell migration and cell stretching can all cause DNA damage^32, 33^. Interestingly, we observed an enrichment for genes involved in cytoskeletal changes but not oxidative stress in our pathway analysis. Since, AEC2 differentiation into AEC1 requires extensive cell stretching and cytoskeletal dynamics (Fig. 4) we hypothesized that PATS cells experience significant mechanical stretching that can lead to DNA damage. To test this hypothesis, we purified AEC2s from *SFTPC-GFP* mice and cultured them on a plastic surface (2D cultures), conditions under which AEC2s stretch and differentiate into AEC1s ^34^. Within 5 days after plating, most AEC2s stretched and either lost (GFP^-^) or downregulated (GFP^lo^) GFP expression. These GFP^-^ and GFP^lo^ cells increased their surface area through stretching and spreading and began to express AGER, a marker for AEC1. Interestingly, these AGER+ cells are also positive for the DNA damage marker, γH2AX (Fig. 5i,k). Of note, we did not see cell stretching and DNA damage markers in cells that we collected on day-2 after plating (Fig. 5k). Taken together, our data show that cuboidal AEC2s that differentiate into large and thin AEC1 during alveolar regeneration naturally experience DNA damage and repair and undergo transient senescence.

### Ectopic activation of TP53 signaling promotes AEC2 to AEC1 differentiation via PATS during alveolar injury-repair

Our data suggest that activation of TP53 signaling is associated with PATS during AEC2 differentiation into AEC1 after lung injury (Fig. 4). Furthermore, previous studies in other tissues have suggested that TP53 signaling promotes differentiation and suppresses stem cell self-renewal ^35^. This led us to hypothesize that enhanced activation of TP53 signaling increases AEC2 differentiation into AEC1 involving PATS after bleomycin-induced lung injury. To test this hypothesis, we utilized the *Sftpc-CreER;R26-tdTomato* mouse model, in which tamoxifen administration is followed by bleomycin or PBS. These mice were then administered Nutlin-3a (an activator of the TP53 pathway) or DMSO (control) daily starting on day-8 after bleomycin and tissues were collected on day-18 (Fig. 6a). Immunostaining analysis for the AEC1 cell marker, AGER, revealed that Nultin-3a treatment led to significantly greater differentiation of lineage labelled AEC2 cells into AEC1 compared to DMSO controls (Fig. 6b). We did not observe any AGER^+^*Sftpc*-tdt^+^ cells in uninjured lungs that received Nutlin-3a, suggesting that in the absence of injury, Nutlin3a-induced TP53 activation alone is not sufficient to induce AEC2 differentiation into AEC1 (Fig. 6b). Our quantitative analysis revealed that 67.25%±1.94% AGER^+^ cells are *Sftpc*-tdt^+^ after bleomycin injury and Nutlin-3a treatment compare to 54.55%±3.71% in bleomycin and DMSO treated animals. Additionally, we also tested the effect of Nutlin-3a on AEC2s in alveolar organoid cultures. Organoid cultures were treated with Nutlin-3a or DMSO starting on day-7 until harvest on day 15 (Fig. 6d). Immunostaining analysis revealed that organoids treated with Nutlin-3a showed both an increase in the number and the expression intensity of AGER^+^ cells and a decrease in the number of proliferating cells as demonstrated by Ki67 immunostaining compare to controls (Fig. 6d,e). These data further support our hypothesis that TP53 signaling enhances AEC2 differentiation both *in vivo* regenerating tissues and *ex vivo* organoid cultures.

**Figure 6.**
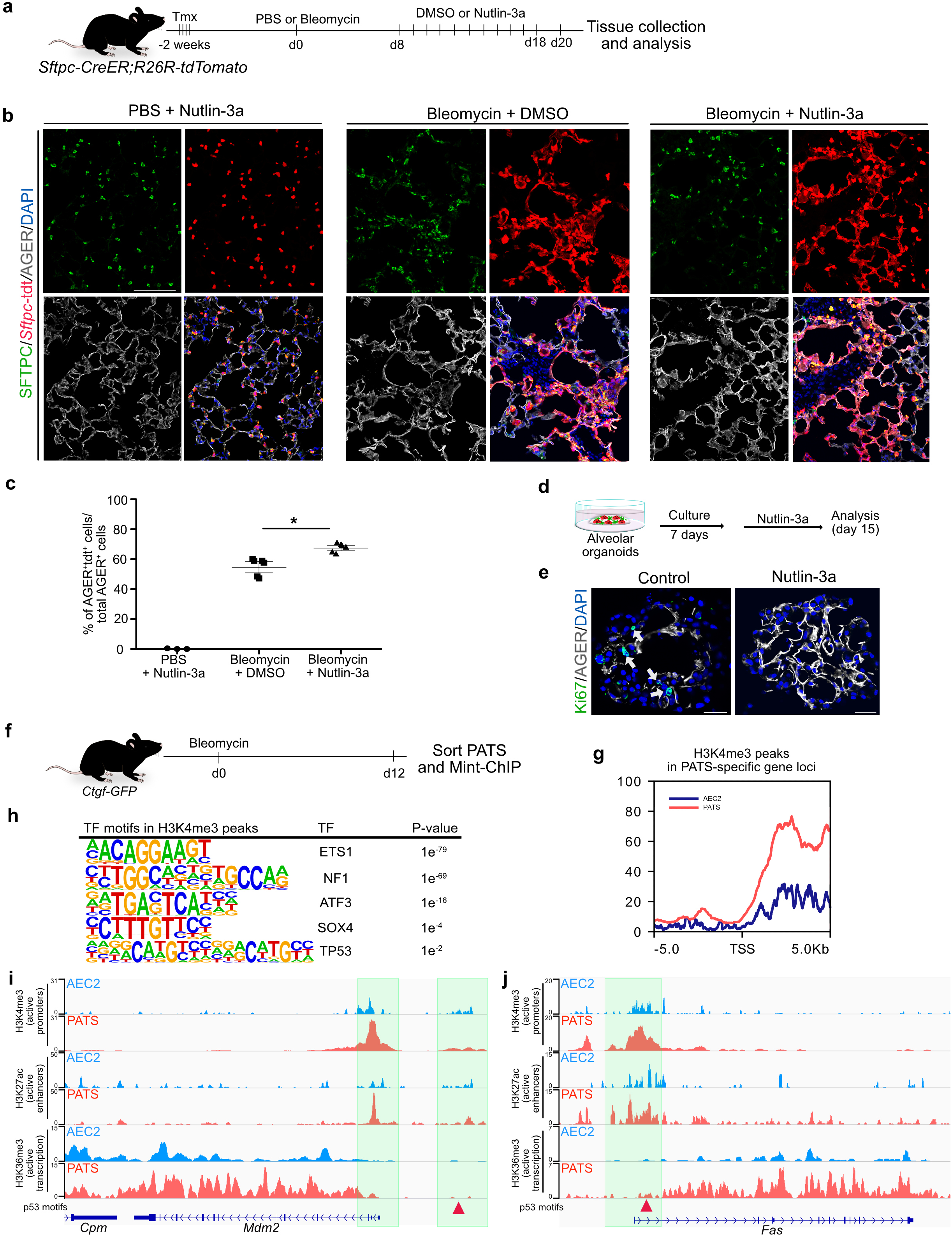
Transcriptional control of PATS by TP53 signaling. a. Experimental workflow for sequential administration of tamoxifen followed by PBS or bleomycin (d0) administration and Nutlin-3a or DMSO treatment (d8-18) and tissue collection (d20) for analysis using *Sftpc-CreER;R26R-tdTomato* mice. b, Co-staining of SFTPC (green), *Sftpc*-tdt (red) and AGER (grey) in PBS+Nutlin-3a (left panel), bleomycin+DMSO (middle panel) and bleomycin+Nutlin-3a (right panel) treated mice. DAPI stains nuclei (blue). Scale bar: 100 µm. c, Quantification of AGER^+^tdt^+^ cells in total AGER^+^ normalized to *Sftpc*-tdt labeling efficiency. Asterisk shows *p* = 0.036. Data are from three independent experiments and are presented as mean ± s.e.m. d. Schematic of alveolar organoid culture treated with Nutlin-3a. e, Immunostaining for Ki67 (green) and AGER (grey) in control or Nutlin-3a treated alveolar organoids. DAPI stains nuclei (blue). Scale bar: 30 µm. f, Schematic of bleomycin-induced lung injury in *Ctgf-GFP* mice. g, Distribution of H3K4me3 peaks in PATS marker gene loci in PATS (red line) and homeostatic AEC2 (blue line). i,j, Transcriptional activity of known TP53 target genes (*Mdm2* and *Fas*) in PATS compared to AEC2s. Arrowhead indicates location of predicted TP53 binding motifs. Green-shade regions are promoter or enhancer.

### Transcriptional control of PATS by transcription factor TP53

To identify the transcription factor modules that are potentially active in PATS, we performed chromatin immunoprecipitation analysis for Histone H3 lysine 4 tri-methylation (H3K4me3), Histone H3 lysine 36 tri-methylation (H3K36me3), and Histone H3 lysine 27 acetylation (H3K27ac), to identify active promoters, transcribing genes, and enhancers, respectively. We isolated and sorted PATS from *Ctgf-GFP* mice after bleomycin administration and AEC2s from control mice. Sorted cells were used for profiling the above-mentioned histone marks using Mint-ChIP, a recently described method that requires fewer cell numbers (Fig. 6f) ^36^. As expected, we found H3K4me3 enrichment in genes corresponding to cell type specific promoters (for example, *Sftpc in* AEC2s and *Fn1* in PATS) (Supplementary Fig. 7a,b). Further analysis revealed enrichment of numerous H3K4me3 peaks overlapping with transcriptional start sites (TSS) and promoters of transcripts specific to PATS from bleomycin exposed lungs compared to AEC2s from controls (Fig. 6g-h). Motif analysis was performed to predict enrichment of binding sites for transcription factors (TFs) in H3K4me3 and H3K27ac peaks. We found enrichment of binding sites for TFs including TP53, ETS1, NF1, ATF3, and SOX4, all of which have been implicated in PATS enriched pathways such as cell cycle arrest, senescence, DNA damage repair, and cytoskeletal control (Fig. 6g,h) ^37–40^. For example, we found predicted binding sites for TP53 in the *Mdm2* enhancer and *Fas* promoter, two well-known direct targets of TP53 (Fig. 6i,j) ^41^. Similarly, we found predicted binding sites for NF1, ETS1, and ATF3 in the promoter regions of PATS specific genes including *Lgals3 and Sox4* (Supplementary Fig. 7c,d). Taken together, our data implicate a direct role for TP53 in transcriptional control of PATS-specific genes.

### PATS associated gene expression signatures and signaling pathways are enriched in human fibrotic lungs

Numerous studies have suggested that abnormal ECM remodeling and chronic inflammation and stress pathways are associated with defective regeneration in multiple organs, including the lung ^42–45^. Indeed, recent studies have suggested that alveolar epithelial cells that line fibrotic foci in idiopathic pulmonary fibrosis (IPF) show features of senescence, growth arrest, and differentiation blockade ^46–48^. Thus, we hypothesized that in response to a non-permissive pathologic microenvironment, alveolar progenitors stall their differentiation process in a PATS-like state. To address this hypothesis, first we analyzed a recently reported scRNA-seq dataset from human IPF lungs ^49^. First, we segregated alveolar epithelial cells by excluding all non-epithelial cells and airway cells from further analysis. We observed numerous AEC2s and AEC1s in both healthy and IPF lungs, with apparent overlap in UMAP plots (Fig. 7a-c). By contrast, we found a distinct cell cluster that is highly enriched in IPF samples and did not overlap with either AEC1 or AEC2 (Fig. 7a,b). Previously, these cells were annotated by their marker gene expression as KRT5^-^/KRT17^+49^. Significantly, differential gene expression analysis revealed a striking resemblance of transcripts, including *Sfn, Sox4* and *Fn1,* between the IPF-enriched cell cluster and PATS from our *ex vivo* organoids and LPS-induced murine lung injury model *in vivo* (Fig. 1, Fig. 4, Fig. 7a-e, and Supplementary Fig. 8a,b). We therefore named this human IPF-specific cluster as ‘PATS-like’ cells (Fig. 7a-c). Other transcripts that are highly enriched in the PATS-like population are *KRT17, CALD1, PRSS2, MMP7,* and *S100A2.* Gene ontology and pathway analysis revealed an enrichment for components of p53 signaling (*CDKN1A, CDKN2A*, and *MDM2*), DNA-damage checkpoint (*RPS27L* and *PLK3*) and cellular senescence (*TGFB2 and HIPK2*) (Fig. 7d-f and Supplementary Fig. 8). Moreover, the PATS-like cluster is enriched for components of focal adhesion, tight junction, and regulation of actin cytoskeleton, indicating a remarkable resemblance between PATS-like cells in IPF and those in organoids and regenerating alveoli *in vivo* (Fig. 4 and Supplementary Fig. 8c). We also found a significant enrichment for *AREG*, *TGFB1*, *TGFB2* and *TIMP1*, known regulators of fibrosis, in PATS-like cells (Fig. 7f). To validate single cell transcriptome data, we performed immunofluorescence analysis on human lung sections from healthy and IPF samples (Supplementary Fig. 9a). Co-immunofluorescence analysis for markers of PATS-like cells (SFN, CLDN4, LGALS3 and KRT17), AEC2 (HTII-280), AEC1 (AGER) and myofibroblasts (ACTA2) revealed the expression of PATS-like markers specifically in IPF lungs but not in healthy controls and healthy appearing regions in IPF lungs (Fig. 7g-i and Supplementary Fig. 9). Interestingly, KRT17 has been recently shown to be expressed in basal cells of the normal lung and basaloid-like cells in pulmonary fibrotic lungs ^49, 50^. In addition, we also found similar expression pattern of TP63, another marker for basal cells (Supplementary Fig. 9). We find that markers of PATS-like cells are almost exclusively present in fibrotic regions as assessed based on high levels of ECM deposition and accumulation of myofibroblasts (Fig. 7g-i and Supplementary Fig. 9). Quantitative analysis for SFN^+^ cells in total HTII-280^+^ cells further supported these findings (91.07%±3% in IPF samples vs 4.16% ±1.02% in healthy human) (Fig. 7h). Moreover, we observed that SFN^+^, HTII-280^+^ cells lost their cuboidal shape and acquired an elongated morphology in IPF lungs as opposed to cuboidal AEC2 or extremely thin and elongated AEC1 in healthy lungs (Fig. 7g,i and Supplementary Fig. 9b,c). Of note, AGER expression is absent in PATS-like cells in IPF lungs (Fig. 7i). Next, we analyzed markers of cell senescence (β-galactosidase activity and CDKN1A/p21) and DNA damage response (γH2AX) in IPF and healthy lungs. Our data revealed specific expression of these markers in the PATS-like population in IPF but not in healthy lungs (Fig. 7 j,k and Supplementary Fig. 9 d,e). Taken together, our analysis revealed similarities between PATS from regenerating tissues and PATS-like cells that are specific to pathological fibrotic lungs.

**Figure 7.**
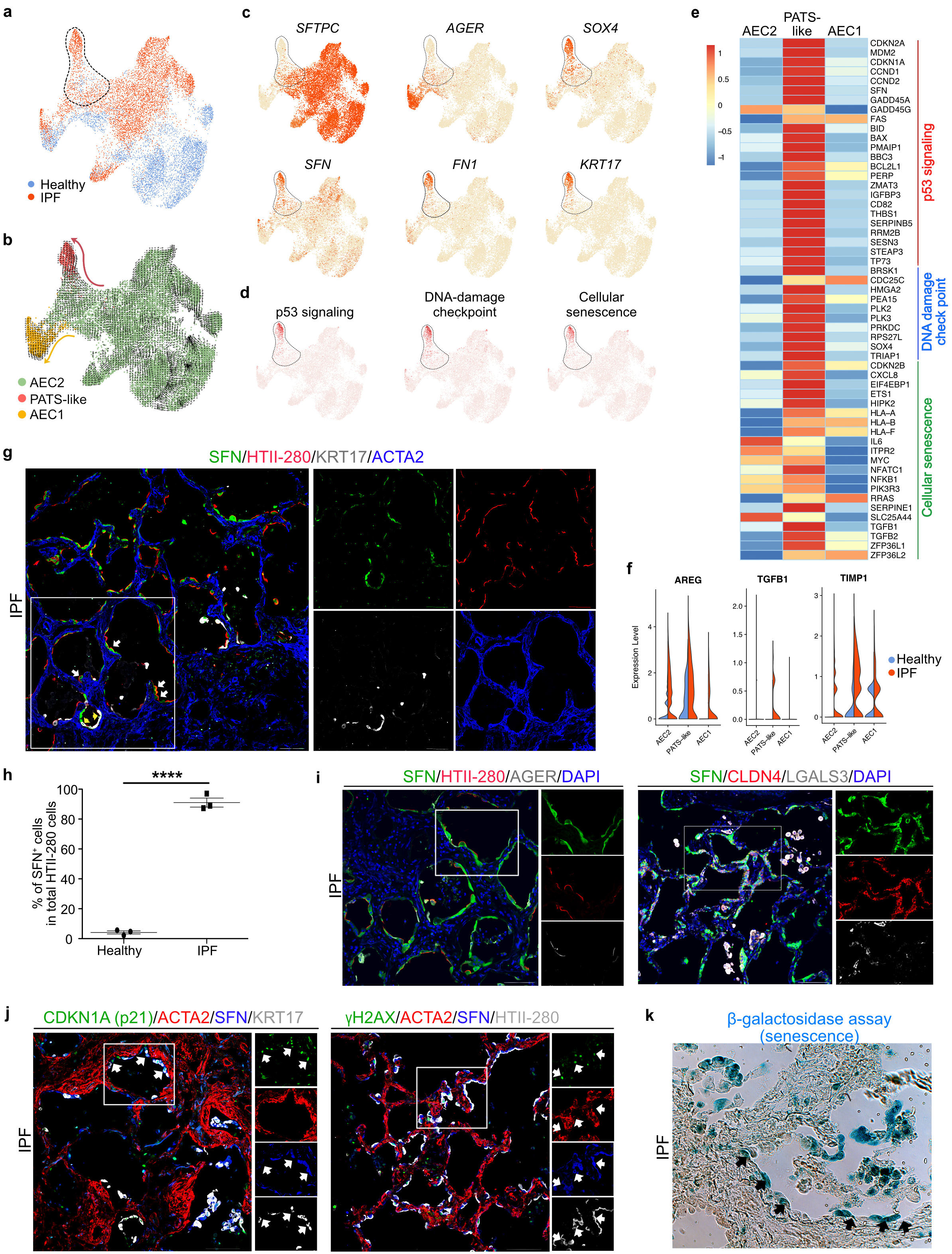
Enrichment of PATS-like states in IPF suggests persistence of this state in pathological milieu. a, UMAP shows scRNA-seq data from alveolar epithelial cells in healthy and IPF lungs. b, RNA velocity analysis predicts lineage trajectories in alveolar epithelial cell populations. Arrows indicates strong RNA velocities. c, UMAP plots show the expression of indicated genes in healthy and IPF lung scRNA-seq data. d, UMAP plots show enrichment of candidate signaling pathways healthy and IPF lung scRNA-seq data. e, Heatmap showing expression of known target genes of indicated signaling pathways in AEC1, AEC2, and PATS-like state. Scale indicates z-score where red is high, and blue is low. f, Violin plots showing IPF-relevant gene expression in indicated cell types/cell states in control and IPF lungs. g-h, Co-staining for PATS-like markers in human IPF lungs. g, Quadruple immunostaining for SFN (green), HTII-280 (red), KRT17 (grey) and ACTA2 (blue) in IPF lung. White arrows indicate SFN^+^, HTII-280^+^ cells. Yellow arrowheads demonstrate SFN^+^, KRT17, HTII-280^+^ cells. h, Quantification of SFN^+^ cells in total HTII-280^+^ cells. Data are from three independent experiments and are presented as mean ± s.e.m. Asterisks indicate *p* < 0.0001. i, Left panel shows triple immunostaining for SFN (green), HTII-280 (red) and AGER (grey) and right panel shows triple immunostaining for SFN (green), CLDN4 (red) and LGALS (grey). j, Quadruple immunostaining for senescence marker p21 (green), in combination with SFN (blue), ACTA2 (red) and KRT17 (grey). White arrows show SFN^+^, KRT17^+^, p21^+^ triple positive cells surrounded by ACTA2 (red) positive cells (left panel). Quadruple immunostaining for γH2AX (green), SFN (blue), HTII-280 (grey) and ACTA2 (red) in IPF lungs (right panel). Inset indicates region of single channel images shown in left side. Scale bars in g-i indicate 100 µm. j, k, β-galactosidase staining in IPF lung. Black arrows indicate X-gal staining in epithelial cells. See Supplementary Fig. 9 for all corresponding immunostainings on control lungs. Scale bar indicates 100 μm.

## Discussion

The work reported here has uncovered a previously unknown transitional state (PATS) that traverses between AEC2 and AEC1 and has a specific gene expression signature in both *ex vivo* organoid cultures and *in vivo* regenerating tissues. These findings further highlight the resemblance between *ex vivo* organoid models and their *in vivo* correlates. Our fate mapping studies using *Sftpc*-driven and *Krt19*-driven CreER alleles demonstrate a clear linear lineage relationship between AEC2 to PATS to AEC1. Therefore, with the addition of this novel transitional cell state, our study revises the alveolar epithelial cell hierarchy. Based on distinct gene expression signatures, our study indicates that PATS cells are not merely undergoing a gradual loss of AEC2 characteristics but represent a unique transitional population. These cells are identified by their expression of many pathways, including, TP53, NF-κB, YAP, TGFβ, and HIF1, previously shown to be important for lung regeneration ^20, 22, 30, 51, 52^. In addition, we observed enrichment for transcripts associated with cell cycle arrest, senescence TP53 and TGFβ signaling, and the transcription factor SOX4, a known regulator of epithelial-mesenchymal transition and cell adhesion in other tissues ^53–55^. These transcriptional signatures are consistent with the dramatic morphological changes that occur during AEC2 to AEC1 differentiation. Through pharmacological modulation, our data demonstrate that TP53 signaling promotes AEC2 differentiation into AEC1 involving PATS in both *ex vivo* organoid cultures and *in vivo* regenerating tissues. Our analysis of human lungs identified PATS-like cells that are specifically present in alveoli of lungs with pulmonary fibrosis. Similar to murine cells, PATS-like cells in human lungs are also characterized by enrichment for genes associated with cellular senescence and TP53 signaling, as well as TGFβ regulated genes, all of which have been implicated in the pathogenesis of fibrosis multiple organs, including the lung ^56, 57^. In contrast, we also found some differences in gene expression signatures between murine PATS and human IPF-specific PATS-like cells. Some of them include, TP63, KRT17, and COL1A1, which are found only in human PATS-like cells. Interestingly, these genes are characteristic markers of basal cells of the normal airway. Our RNA velocity projections from human scRNA-seq data suggest that these KRt17^+^/TP63^+^ cells originate from AEC2. Indeed, our immunofluorescence studies for KRT17 and TP63 further suggested that these cells are surrounded by cells that express markers of PATS within the same alveoli. All together, these data suggest that KRT17^+^/TP63^+^ cells originate from AEC2.

Another significant finding from this study is that PATS cells undergo extensive stretching and spreading, which makes them vulnerable to DNA damage, a feature commonly associated with most degenerative lung diseases, notably pulmonary fibrosis and cancers ^58–62^. Of note, we did not find cells with DNA damage in AEC1 of lungs that had fully regenerated following bleomycin-induced injury, suggesting that alveolar cells can efficiently repair the damage DNA (data not shown). However, it is also possible that tissue may eliminate such “unfit cells” (with DNA damage) through cell extrusion or cell death mechanisms. Nevertheless, our study uncovered a novel transitional cell state that is vulnerable to DNA damage during AEC2 differentiation into AEC1. Previous studies have revealed that cell stretching causes DNA damage when they migrate or squeeze through narrow spaces ^63^. Indeed, our *in vivo* AEC1 ablation and *ex vivo* 2D culture models suggest similar mechanisms are at play when cuboidal AEC2 undergo extensive stretching during their differentiation into thin and large AEC1. Therefore, the novel transitional cell state has clinical implications as most degenerative lung diseases are accompanied by the presence of cells with DNA damage accumulation ^59, 60^. Interestingly, the PATS population is enriched for SOX4, a transcription factor known to regulate cytoskeletal genes, which is induced following DNA damage and has a crucial role in TP53 stabilization and function^40^. Altogether, these data support a model in which, cells evolved co-transcriptional programs to combat DNA damage that can occur when cells undergo extreme stretching. In addition, genome-wide association studies have identified mutations in DNA damage repair pathways components, such as *XRCC* family genes, *LIG4*, *TERC*, PARP, and *RTEL1* with emphysema and pulmonary fibrosis^58, 64^. Thus, the novel transitional state we have identified here implicates cell shape changes and associated vulnerabilities accompanying alveolar stem cell differentiation in the pathogenesis of some lung diseases (Fig. 8).

**Figure 8.**
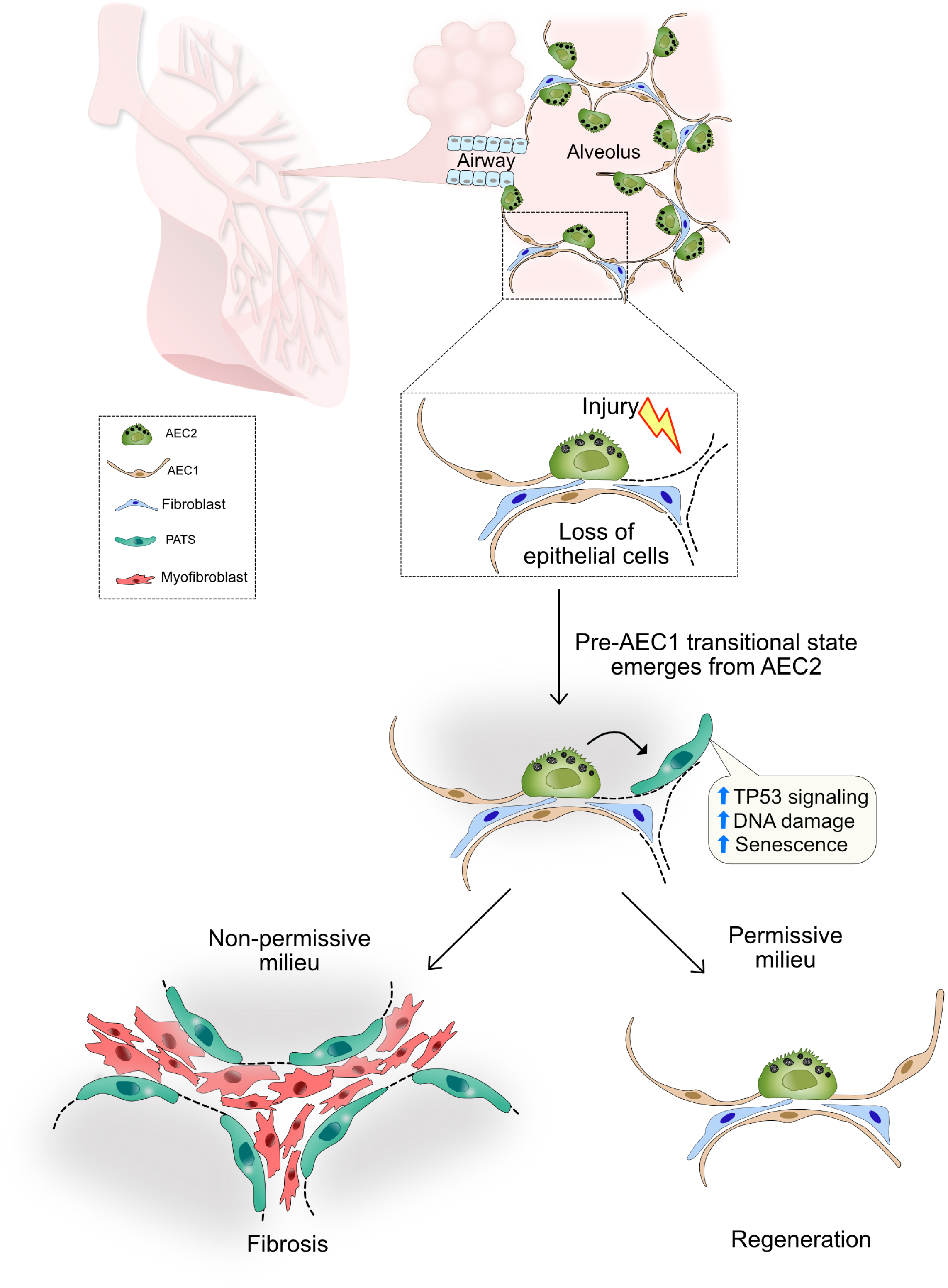
Schematic describing alveolar stem cell-mediated epithelial regeneration and disease pathogenesis. Alveolar stem cells replicate in response to damage and generate a novel transitional cell state which normally matures into functional alveolar type-1 epithelial cells. The newly identified transitional state is vulnerable to DNA damage and undergoes a transient senescent state. This novel state is enriched in human fibrotic lungs.

Senescence is often seen as an age-associated pathological state in which cells acquire an abnormal and irreversible state^48, 62, 65, 66^. Here, we find that alveolar stem cell differentiation involves a novel transitional state which exhibits cardinal features of senescence in normal tissue regeneration. Indeed, prior studies have found senescent cells in embryos in developing limb bud tissues^67^. Therefore, we propose that senescence may not necessarily occur exclusively in aged tissues but can be a reversible transient state accompanying tissue repair or regeneration. Our study thus redefines senescence as a state that can occur as part of normal tissue maintenance programs and can be derailed in human diseases.

In conclusion, using alveolar organoid cultures and *in vivo* injury-repair models, we have identified a novel pre-AEC1 transitional state in lung regeneration. This novel and unique state is associated with cellular senescence and enrichment for pathways known to be associated with defective alveolar regeneration. These results strongly suggest that prolonged senescence and stress mediated pathways in transitional cell states can lead to diseases such as fibrosis.

## Online Methods

### Mice

*Sftpc^tm1(cre/ERT2)Blh^* (*Sftpc-CreER*) ^25^, *Krt19^tm1(cre/ERT)Ggu^/J* (*Krt19-CreER*) ^68^, *Rosa26R-CAG-lsl-tdTomato* ^69^ (crossed with *Sftpc-CreER*), *B6.Cg-Gt(ROSA)26Sor^tm14(CAG-tdTomato)Hze^/J* (*R26R-tdTomato*) ^70^ (crossed with *Krt19-CreER*), *Tg(SFTPC-GFP)#Heat (Sftpc-GFP)* ^71^, *B6.Cg-Ager^tm2.1(cre/ERT2)Blh^/J* (*Ager-CreER*) ^72^, *C57BL/6-Gt(ROSA)26Sor^tm1(HBEGF)Awai^/J* (*R26R-DTR*) ^73^, *Mki67^tm1.1Cle^/J* (*Mki67RFP*) ^74^ and *Tg(Ctgf-EGFP)FX156Gsat* (*Ctgf-GFP*) ^23^ were utilized for experiments. For lineage tracing with *Sftpc-CreER;R26R-tdTomato*, 0.2 mg/g body weight Tamoxifen (Tmx, Sigma-Aldrich) was given via oral gavage or intraperitoneal injection. For lineage tracing using *Krt19-CreER;R26R-tdTomato*, one dose of 0.1 mg/g body weight tamoxifen was given via intraperitoneal injection seven days after bleomycin injury or PBS administration. Animal experiments were approved by the Duke University Institutional Animal Care and Use Committee.

### Bleomycin injury

For bleomycin-induced lung injury, 2.5 U/Kg bleomycin was administered intranasally at two weeks after tamoxifen injection and mice were monitored daily. PBS administered mice served as controls. Mice were sacrificed at different time points after bleomycin injury.

### Diphtheria toxin (DT) administration

Two weeks before DT administration, *Ager-CreER;R26R-DTR* received tamoxifen via IP injection. One dose of 3 µg diphtheria toxin (Millipore #322326) was administer via intraperitoneal injection and mice were sacrificed six days later for tissue collection and analysis.

### Mouse lung dissociation and fluorescence assisted cell sorting

Lung dissociation and FACS were performed as described previously^19^. Briefly, lungs were intratracheally inflated with 1 ml of enzyme solution (Dispase (5 U/ml, Corning #354235), DNase I (0.33 U/ml) and collagenase type I (450 U/ml, Gibco #17100-017)) in DMEM/F12. Separated lung lobes were diced and incubated with 3 ml enzyme solution for 25 min at 37°C with rotation. The reaction was quenched with an equal amount of medium containing 10% Fetal bovine serum (FBS) and filtered through a 100 µm strainer.

The cell pellet was resuspended in red blood cell lysis buffer (155 mM NH_4_Cl, 12 mM NaHCO_3_, 0.1 mM EDTA) and incubated for 2 min, then filtered through a 40 µm strainer. The cell pellet was resuspended in DMEM/F12 + 2% BSA and stained with following antibodies: EpCAM (eBioscience, G8.8), PDGFRα (BioLegend, APA5) and Lysotracker (Thermo Fisher, L7526) as described previously^29^. Sorting was performed using a BD FACS Vantage SE, SONY SH800S or Beckman Coulter MoFlo Astrios EQ.

### Alveolar organoid culture

Alveolar organoid culture was performed as described previously ^10^. Briefly, lineage labeled AEC2s (1-3 × 10^3^) from *SFTPC-GFP* or *Sftpc-CreER;R26R-lsl-tdTomato* mice treated with Tmx were FACS-sorted and PDGFRα^+^ (5 × 10^4^) fibroblasts were resuspended in MTEC/Plus and mixed with equal amount of growth factor-reduced Matrigel (Corning #354230). Medium was changed every other day.

### Nutlin-3a treatment

For *in vivo* studies*, Sftpc-CreER;R26R-tdTomato* were injected with one dose of tamoxifen and rested for two weeks followed by bleomycin or PBS administration. Eight days after injury, Nutlin-3a (Selleckchem #S8059) or DMSO (control) was administered by intraperitoneal injection at concentration of 20mg/Kg/day for ten consecutive days and samples were collected on 20 days after bleomycin administration. For *Ex vivo* study, alveolar organoids were grown for seven days followed by Nutlin-3a (2 µM) treatment for 8 days before their harvest.

### Droplet-based single-cell RNA sequencing (Drop-seq)

Organoids embedded in Matrigel were incubated with *Accutase* solution (sigma #A6964) at 37°C for 20 min followed by incubation with 0.25% Trypsin-EDTA at 37°C for 10 min. Trypsin was inactivated using DMEM/F-12 Ham supplemented with 10% FBS and cells were then resuspended in PBS supplemented with 0.01% BSA. After filtration through 40 µm strainer, cells at a concentration of 100 cells/µl were run through microfluidic channels at 3,000 µl/h, together with mRNA capture beads at 3,000 µl/h and droplet-generation oil at 13,000 µl/h. DNA polymerase for pre-amplification step (1 cycle of 95°C for 3 min, 15-17 cycles of 98°C for 15 sec, 65°C for 30 sec, 68°C for 4 min and 1 cycle of 72°C for 10 min) was replaced by Terra PCR Direct Polymerase (#639271, Takara). The other processes were performed as described in the original Drop-seq protocol ^75^. Libraries were sequenced using HiSeq X with 150-bp paired end sequencing.

### Computational analysis for scRNA-seq

scRNA-seq analysis of alveolar organoids was performed by processing FASTQ files using dropSeqPipe v0.3 (https://hoohm.github.io/dropSeqPipe) and mapped on the GRCm38 genome reference with annotation version 91. Unique molecular identifier (UMI) counts were then further analyzed using an R package Seurat v3.1.1 ^76^. UMI count matrix of murine lungs treated with LPS (GSE130148) ^22^ was obtained from Gene Expression Omnibus (GEO). UMI counts were normalized using SCTransform. Cell barcodes for the clusters of interests were extracted and utilized for *velocyto run* command in velocyto.py v0.17.15 ^77^ as well as generating RNA velocity plots using velocyto.R v0.6 in combination with an R package SeuratWrappers v0.1.0 (https://github.com/satijalab/seurat-wrappers). Twenty-five nearest neighbors in slope calculation smoothing was used for *RunVelocity* command. After excluding duplets, specific cell clusters were isolated based on enrichment for *Sftpc, S*ftpa1*, S*ftpa2*, Sftpb, Lamp3, Abca3, Hopx, Ager, Akap5, Epcam, Cdh1, Krt7, Krt8, Krt18, Krt19, Scgb1a1* and *Scgb3a1* as well as negative expressions of *Vim, Acta2, Pdgfra* and *Pdgfrb* in UMAP plots. The Rds files for control and idiopathic pulmonary fibrosis (IPF) lungs were obtained from GEO (#GSE135893)^49^. Cell clusters of AEC2, AEC1, transitional AEC2 and *KRT5*^-^/*KRT17*^+^ were extracted and analyzed. Markers for each cluster (Supplementary Table 1) obtained using *FindAllMarkers* command in Seurat were utilized for identifying specific signaling pathways and gene ontology through Enrichr ^78^. Z-scores were calculated based on combined score in Kyoto encyclopedia of genes and genomes (KEGG) to compare enrichment of signaling and ontology across different cell clusters. The results were displayed in heatmap format generated using an R package pheatmap v1.0.12. Scaled data in Seurat object were extracted and mean values of scaled score of gene members in each pathway were calculated and shown in UMAP as enrichment of signaling pathways. The gene member lists of utilized pathways were obtained from KEGG pathways ^79^ and AmiGO ^80^. Log_2_ fold changes and P-values in each gene extracted using *FindMarkers* command in Seurat with Wilcoxon rank sum test were shown in a volcano plot using an R package EnhancedVolcano v1.3.1 (https://github.com/kevinblighe/EnhancedVolcano) to show specific markers for *Ctgf*^+^ cells and *Lgals3*^+^ cells.

### Mint-ChIP (Multiplexed indexed T7 ChIP-seq)

FACS-sorted CD31^-^/CD45^-^/CD140a^-^/CD326^+^/*CTGF*-GFP^+^ cells (PATS) from *Ctgf-GFP* mice treated with Bleomycin (d12) and CD31^-^/CD45^-^/CD326^+^/Lysotracker^+^/*Mki67*-RFP^-^ cells (AEC2) from *Mki67-RFP* homeostatic mice were processed using Mint-ChIP described previously^36^. Open-sourced newest protocol named Mint-ChIP3 in protocol.io was used. Following cell lysis, chromatin was digested with 300 units reaction of MNase (#M0247S, New England Biolabs) at 37°C for 10 min. T7 adapter ligation was performed for 2 hrs and then the samples were split to have ∼7,000 cells per antibody. The samples were incubated with Histone H3 (H3) antibody (1 ul, #39763, Active Motif), Histone H3 lysine 36 trimethylation (H3K36me3) antibody (1 ul, #61101, Active Motif) or Histone H3 lysine 4 trimethylation (H3K4me3) antibody (1 ul, #ab8580, Abcam) and Histone H3 lysine 27 acetylation (H3K27ac) antibody (1 ul, #39133, Active Motif) at 4°C overnight. DNA was purified followed by T7-driving *in vitro* transcription at 37°C for 3 hrs. Reverse transcription was performed as described in original protocol followed by library preparation using Terra Direct PCR polymerase (#639271, TaKaRa). Two experimental replications were performed for each cell type. Libraries were sequenced using Hiseq X or NovaSeq 6000 with at least 5M reads of 150-bp paired end per sample.

### Computational analyses for Mint-ChIP

FASTQ files were generated using Bcl2fastq. Additional demultiplexing for Mint-ChIP FASTQ files were performed using Je ^81^. Low quality reads were trimmed out from FASTQ files using trimmomatic v0.38 ^82^. Reads were mapped on mm10 genome reference using BWA ^83^. HOMER ^84^ was used for generating bedGraph files to visualize them in Integrative Genomics Viewer (IGV)^85^. Peak calling for H3K4me3 was performed using HOMER’s *getDifferentialPeaksReplicates.pl -region -size 1000 -minDist 2000 -C 0 -L 50* with normalization by H3. Motif analysis was performed using HOMER’s *findMotifsGenome.pl*. The packages were run through a pipeline called MintChIP (https://github.com/jianhong/MintChIP). deepTools ^86^ was used for generating a chart of called peaks of H3K4me3. Called peaks of H3K4me3 in both homeostatic AEC2 and PATS after bleomycin-induced lung injury were merged in Fig. 6g using Affinity Designer. Genes shown in Fig. 4a left panel were used to generate Fig. 6g.

### Human lung tissue

Excised subtransplant-quality human lung tissues without preexisting chronic lung diseases were obtained from the Marsico Lung Institute at the University of North Carolina at Chapel Hill under a University of North Carolina Biomedical Institutional Review Board-approved protocol (#03-1396). Informed consent was obtained from all participants where necessary. Samples of explanted fibrotic human lungs were procured through the BioRepository and Precision Pathology Center at Duke University in accordance with institutional procedures (Duke University Pro00082379 – “Human Lung Stem Cells”; exempt research as described in 45 CFR 46.102(f), 21 CFR 56.102(e) and 21 CFR 812.3(p) which satisfies the Privacy Rule as described in 45CFR164.514). The diagnosis of idiopathic pulmonary fibrosis (IPF) was evaluated by a surgical pathology team. Specimens were washed thoroughly in PBS prior to inflation and immersion in 4% PFA overnight at 4°C. Specimens were subsequently washed in PBS until the appearance of blood was minimal followed by incubation in 30% sucrose at 4°C. Samples were then incubated with 1:1 mixture of OCT for 1 hour at 4°C before embedding in OCT. 7-9 µm thick sections were used for histological analysis.

### Immunostaining

Lungs and alveolar organoids were prepared as described previously. Briefly, tissues were fixed with 4% paraformaldehyde (PFA) at 4°C for 4 h and at room temperature for 30 min then embedded in OCT or Paraffin. 10-µm sectioned samples were utilized for staining following incubation at 95°C for 10-15 min for antigen retrieval using 10 mM sodium citrate. Primary antibodies were as follows: Pro-surfactant protein C (Millipore, ab3786, 1:500), AGER (R&D systems, MAB1179, 1:250), KRT8 (DSHB, TROMA-I, 1:50), KRT17 (NSJ, V2176; 1:250), KRT19 (DSHB, TROMA-III, 1:50), tdTomato (ORIGENE, AB8181-200, 1:500), CLDN4 (Invitrogen, 36-4800, 1:200), GFP (Novos Biologicals, NB100-1770, 1:500), LGALS3 (Cedarlane, CL8942AP, 1:500); SOX4 (Invitrogen, MA5-31424, 1:250), SFN (Invitrogen, PA5-95056, 1:250 or Proteintech, 66251-1-Ig, 1:500), ACTA2 (Sigma, C6198, 1:500), and, gamma-H2AX (R&D, 4418-APC, 1:500).

### β-Galactosidase (X-gal) staining and Hematoxylin & Eosin (H&E) staining

PFA-fixed frozen sections were incubated with X-gal staining buffer containing 1 mg/ml of X-gal (Thermo, R0941), 5 mM K_3_Fe(CN)_6_, 5 mM K_4_Fe(CN)_6_, 2 mM MgCl_2_, 0.01% sodium deocycholate and 0.02% NP-40 at 37°C overnight. Sections were washed 3 time in PBS and mounted. For H&E staining, 10-µm paraffin sections were submerged in Histo-clear and series of ethanol. Mayer’s Hematoxylin was used to stain nuclei, followed by staining using 1% Eosin Y.

### Image acquisition, processing and quantification

Images were captured using Olympus Confocal Microscope FV3000 using a 20×, 40× or 60× objective, a Zeiss wide-field fluorescence microscope (X-gal staining) and a Zeiss Axio Imager Widefield Fluorescence Microscope (H&E). Cells were manually counted based on IHC markers and DAPI. For determination of average intersects per linear distance, a mean linear intercept analysis was conducted as previously described^87^ over the single channel immunofluorescence stain of interest. Images were processed using Olympus CellSens application or ImageJ and figures were prepared using Affinity Designer.

### Statistics

Sample size was not predetermined. Data are presented as means with standard error (s.e.m) to indicate the variation within each experiment. Statistics analysis was performed in GraphPad Prism. A two-tailed Student’s *t*-test was used for the comparison between two experimental conditions. We used Mann Whitney one tailed test for the comparison between two conditions that showed non-normal distributions.

## Acknowledgements

We thank Brigid Hogan for advice and critical reading of the manuscript, Monica Fernandez de Soto for technical support and members of the Tata lab and Barkauskas lab for discussions. We thank Karen Lyons for providing *Ctgf*-GFP mice, Peter van Galen and Yu Xiang for help with Mint-ChIP analysis., the Duke Cancer Institute Flow Cytometry Shared Resource for help with cell sorting, the Duke Sequencing and Genomic Technologies Shared Resource for sequencing NGS-libraries, the Duke University Light Microscopy Core Facility for imaging equipment and consultation, the Duke Biorepository & Precision Pathology Center for providing human specimens, the Duke Compute Cluster for help with sequencing data analysis. The TROMA-I (KRT8) antibody clone developed by Brulet, P/Kemler was obtained from the Developmental Studies Hybridoma Bank, created by the NICHD of the NIH and maintained at The University of Iowa, Department of Biology, Iowa City, IA 52242. Y.K. is a Japan Society for the Promotion of Science Overseas Research Fellow. A.K. is supported by a medical scientist training program fellowship from NHLBI/NIH (F30HL143911). Human scRNA-seq was supported by R01HL145372 (NEB/JAK), Doris Duke Charitable Foundation (JAK), K08HL130595(JAK), and Boehringer Ingelheim Pharmaceuticals (JAK). This work was supported by a Pathways to Independence award from NHLBI/NIH (R00HL127181) to P.R.T. and funds from Regeneration NeXT and Kaganov-MEDx Pulmonary Initiative at Duke University. This work was partially supported by funds from Whitehead foundation and P.R.T. is a Whitehead Scholar.

## Author contributions

Y.K. designed, conceived and performed NGS-related experiments and the computational analysis, and co-wrote the manuscript; A.T. designed, conceived and performed in vivo and ex vivo experiments and immunostaining, and co-wrote the manuscript; A.K. designed and performed AEC1 ablation experiments; H.K. performed organoid experiments. R.F.L. performed histological analysis; J.O. built a pipeline for Mint-ChIP analysis. N.E.B. and J.A.K. provided scRNA-seq data from healthy and IPF human lungs. P.R.T. designed, conceived and supervised the work and co-wrote the manuscript. All authors reviewed and edited the manuscript.

## Additional information

**Extended Data** is available online

### Data availability

The data described in this manuscript are available from the corresponding author upon request. All the NGS sequencing data in this manuscript will be available at NCBI GEO (accession no. GSE will be provided upon acceptance of the manuscript).

## Correspondence and requests for materials

Correspondence and requests for materials should be addressed to P.R.T.

---

**Supplementary Figure 1.**
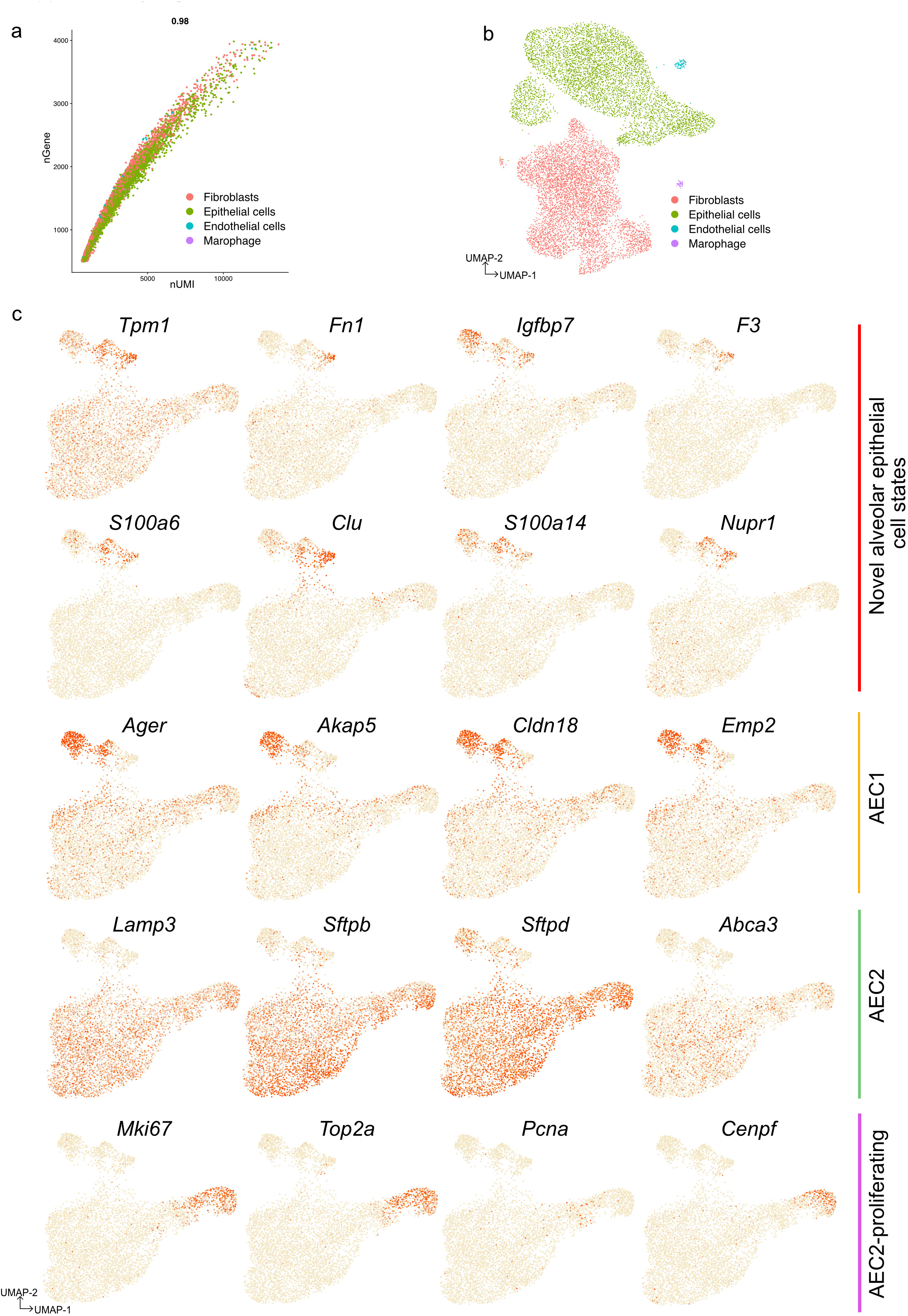
scRNA-seq segregates distinct alveolar population with specific markers. a, Pearson correlation plot visualizes the number of genes per cell (nGene) and unique molecular identifier (nUMI) in total cells derived from alveolar organoids. b, UMAP shows major cell populations including epithelial cells (green), fibroblasts (red), and some minor populations such as endothelial cells (blue) and macrophages (purple) in alveolar organoids. c, UMAP plots show the expression of indicated genes in epithelial cell populations derived from our alveolar organoid scRNA-seq datasets.

**Supplementary Figure 2.**
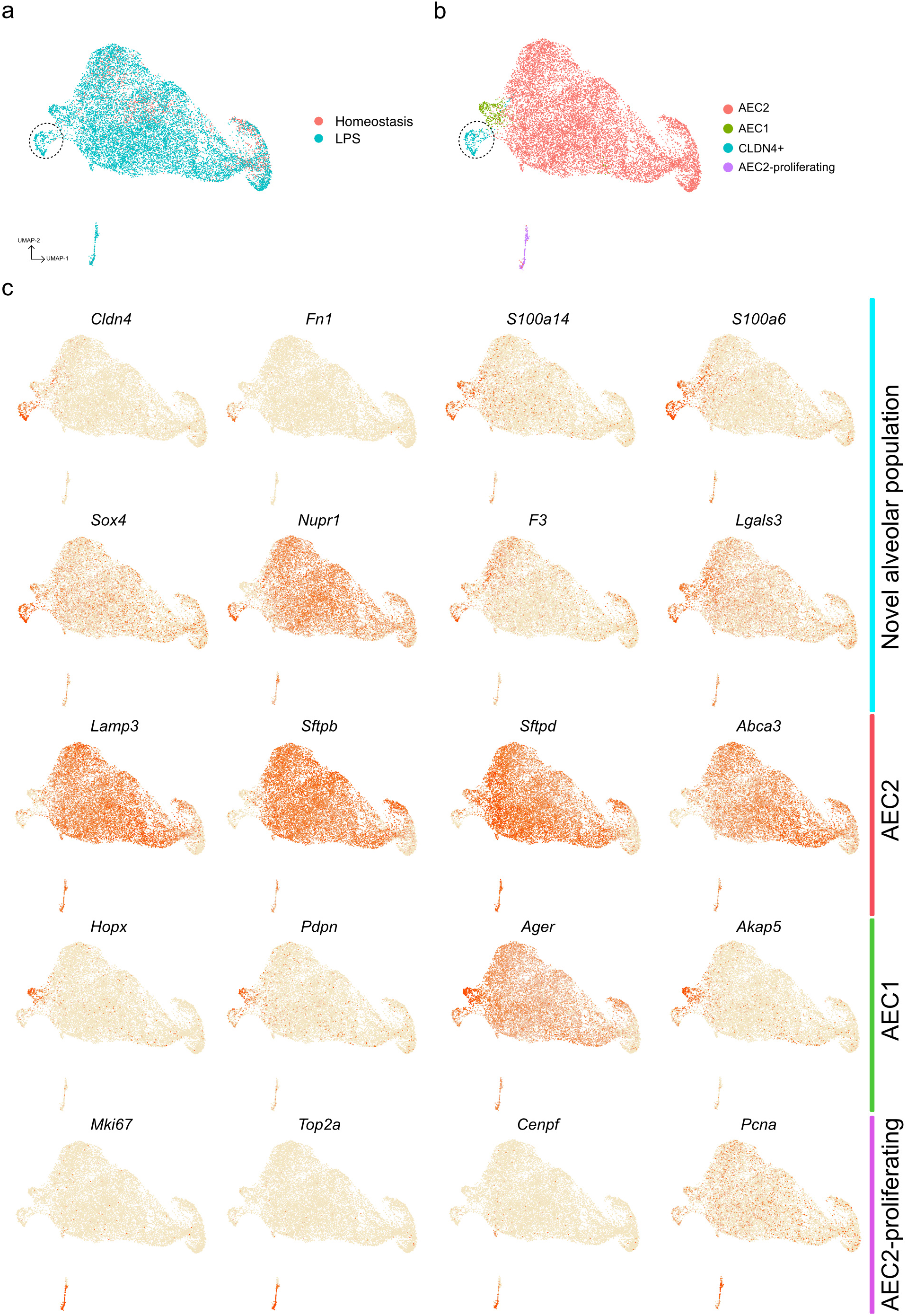
scRNA-seq analysis revealed the emergence of a novel epithelial cell population after LPS-induced lung injury *in vivo*. a, UMAP shows homeostatic (red) and LPS-treated (blue) lung alveolar epithelial cells. b, Distinct cell populations in alveolar epithelial cells in control and LPS-treated lungs. c, UMAP plots show the expression of indicated genes in alveolar epithelial cells in control and LPS-treated lungs.

**Supplementary Figure 3.**
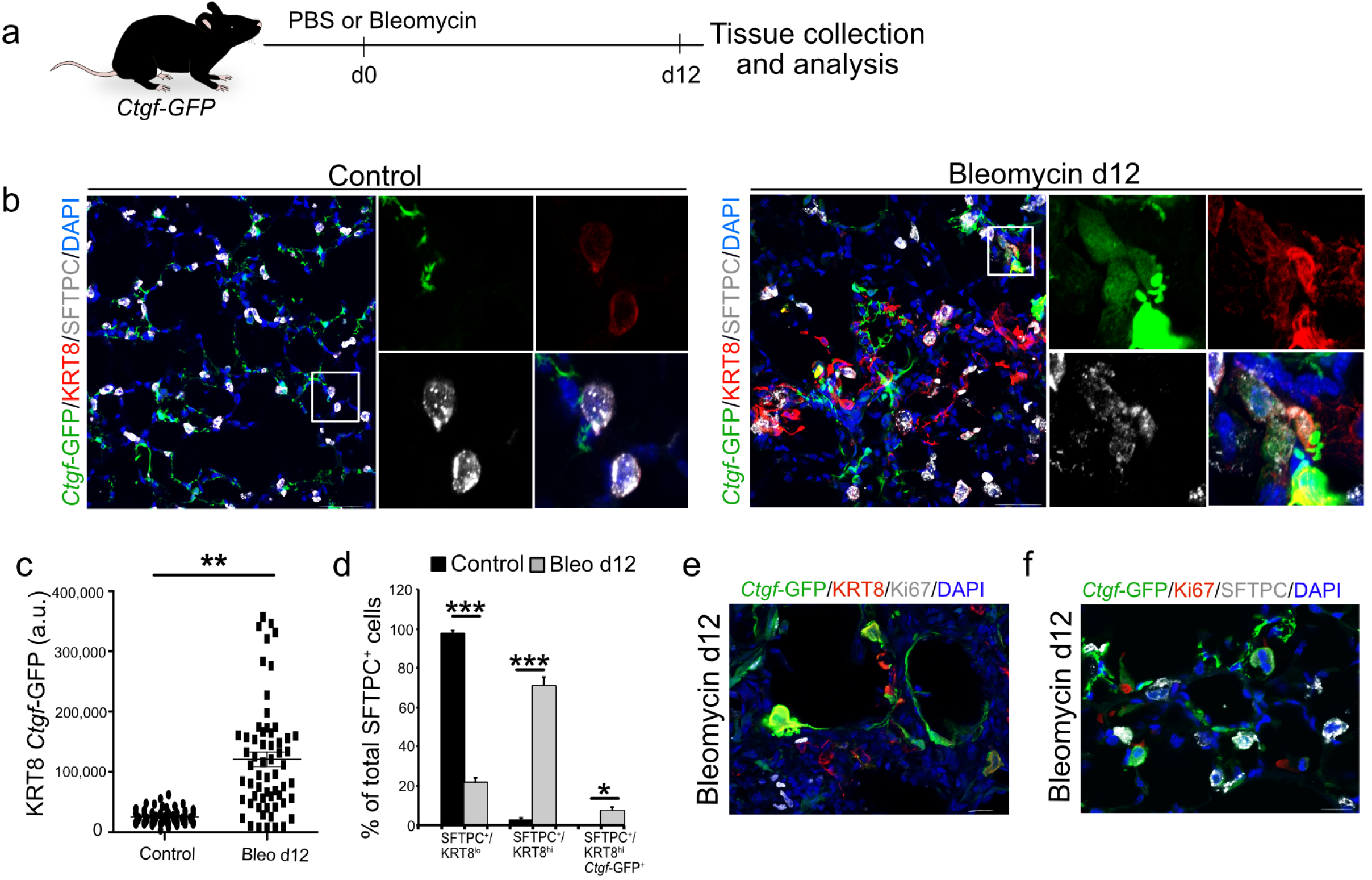
Specific markers are expressed in the novel alveolar population in the lungs following bleomycin treatment *in vivo*. a, Schematic of bleomycin-induced lung injury in *Ctgf*-GFP mice. b, Immunostaining for *Ctgf*-GFP (green), KRT8 (red) and SFTPC (grey) in control lung (left) and bleomycin day 12 injured mice (right). Magnified single channel images are shown on the right. White line box indicates magnified region. c, Quantification of signal intensity of KRT8 immunostaining in control (black circles) and experimental lung 12 days after bleomycin injury (black rectangles); *p* < 0.0001; Wilcoxon rank sum test. d, Quantification of KRT8^lo^, KRT8^hi^ and KRT8^hi^/GFP^+^ subpopulations of SFTPC^+^ cells in control lung (white bars) and lung 12 days after bleomycin injury (gray bars). Error bars, mean ± s.e.m (n = 3). KRT8^lo^- P=1 × 10^-5^; KRT8^hi^- P=1 × 10^-4^; KRT8^hi^ *Ctgf*-GFP^+^- P=2.6 × 10^-3^. e, Immunostaining for *Ctgf*-GFP (green), KRT8 (red) and Ki67 (grey), e, for *Ctgf*-GFP (green), Ki67 (red) and SFTPC (grey) on bleomycin-treated lungs collected on 12 days post injury. DAPI stains nuclei (blue). Scale bars indicate 20 µm.

**Supplementary Figure 4.**
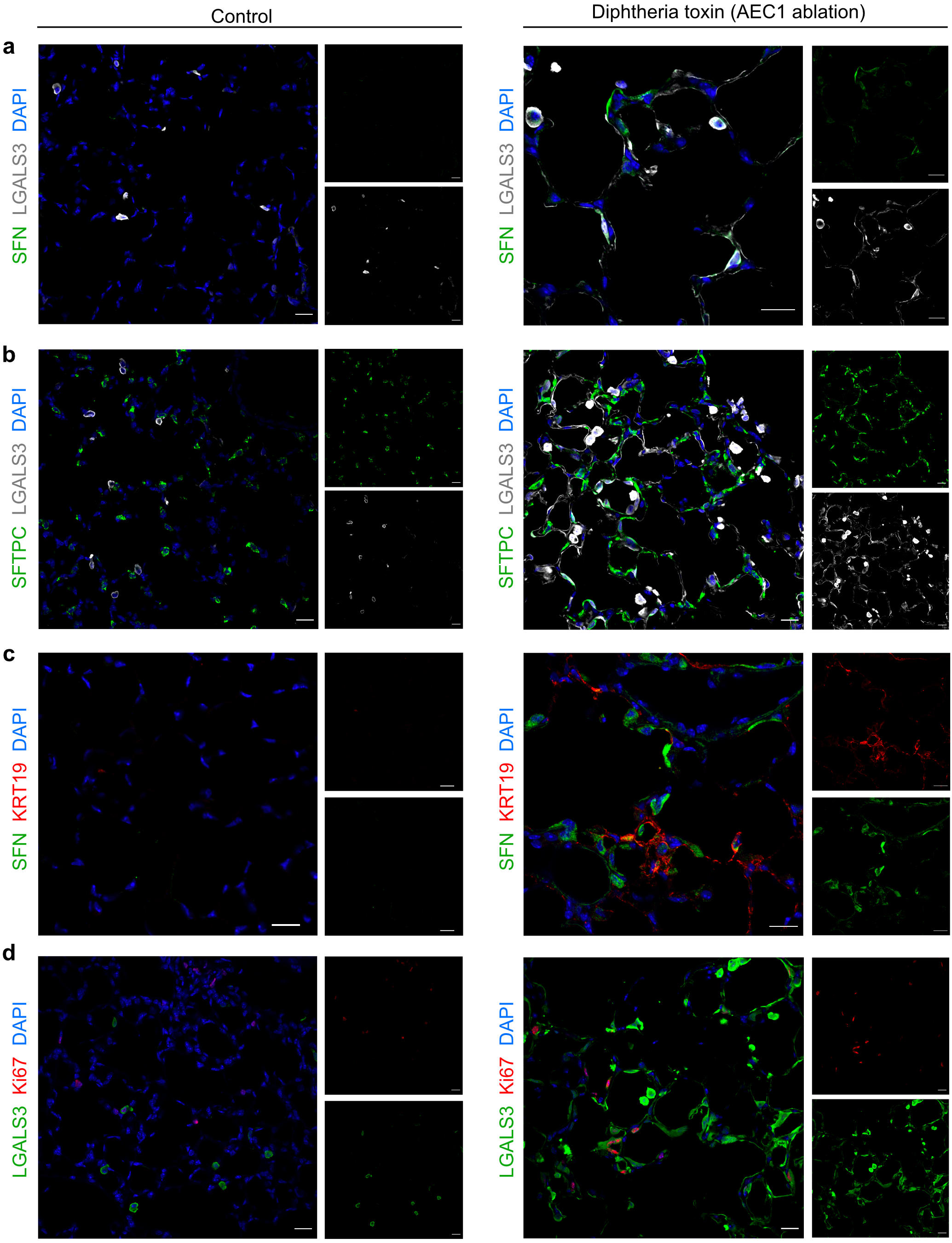
Expression of markers specific to the novel alveolar population in AEC1-specific ablation mouse model. a-d, Co-immunostaining for a) SFN (green) and LGALS3 (grey) or b) SFTPC (green) and LGALS3 (grey) or c) SFN (green) and KRT19 (red) or d) LGALS3 (green) and Ki67 (red) in control (left) and AEC1-ablated lungs (right). DAPI stains nuclei (blue). Scale bars indicate 20 µm.

**Supplementary Figure 5.**
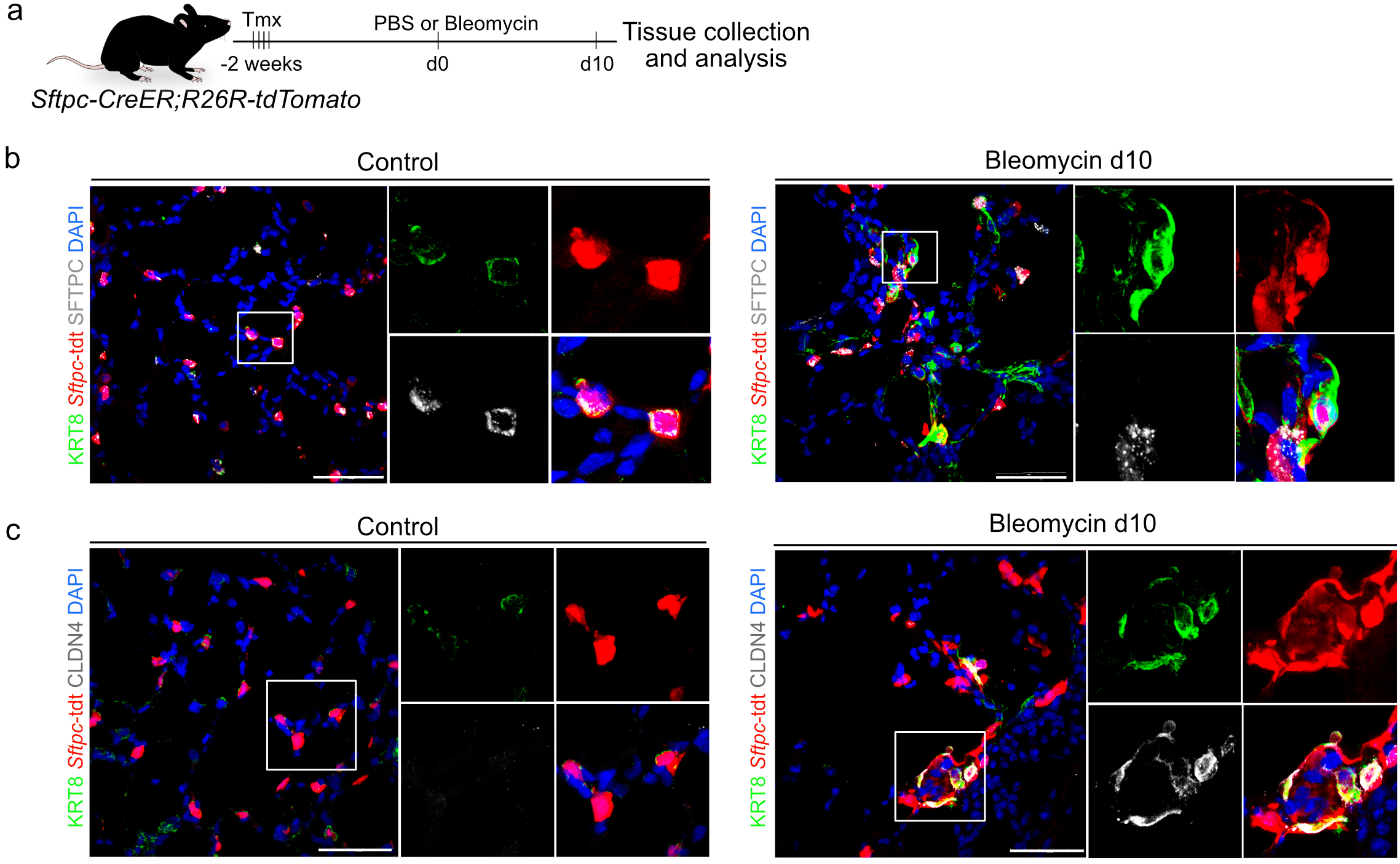
AEC2-derived cells are the cell-of-origin for the novel alveolar cell state following bleomycin-induced lung injury. a, Schematic of AEC2 lineage tracing using *Sftpc-CreER;R26R-tdTomato* mice. Tamoxifen was given 2 weeks prior to PBS or bleomycin administration followed by tissue harvest. b, Immunostaining for KRT8 (green), *Sftpc*-tdt (red) and SFTPC (grey). c, Co-staning for KRT8 (green), *Sftpc*-tdt (red) and CLDN4 (grey) in control (upper) and bleomycin-treated lungs (lower). White boxed insets are shown on the right in each panel. DAPI stains nuclei (blue). Scale bars indicate 50 µm.

**Supplementary Figure 6.**
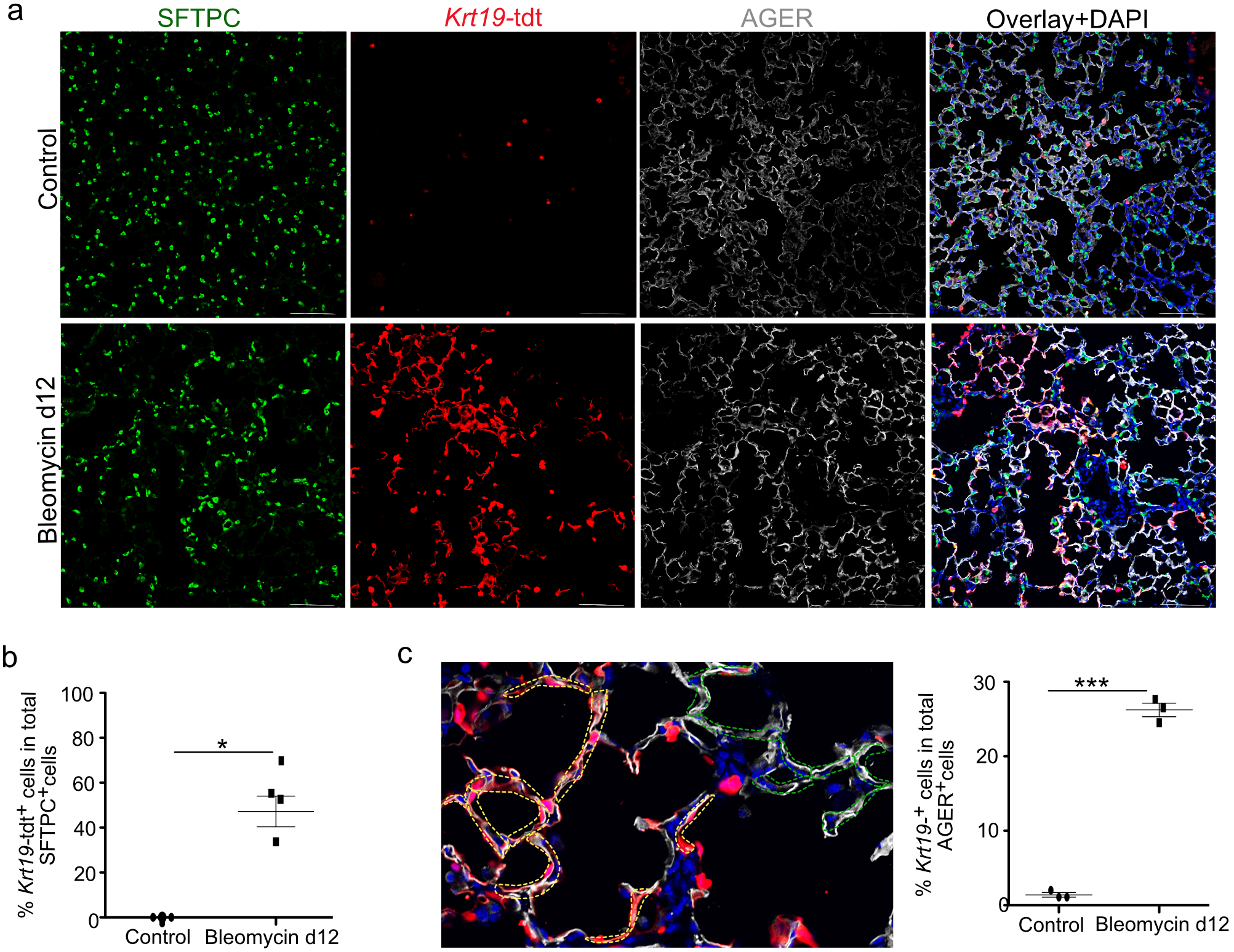
Lineage tracing revealed that PATS differentiate into AEC1 following bleomycin-induced alveolar injury *in vivo*. a, Immunostaining for SFN (green), *Krt19*-tdt (red) and AGER (grey) in control (upper) and bleomycin-treated lungs (lower). Scale bars 100 µm. b, Quantification of *Krt19*-tdt^+^ cells in total SFTPC^+^ cells in injured regions. Asterisk shows *p* < 0.031. (Mann-Whitney test) Data are from three independent experiments and are presented as mean ± s.e.m. c, Quantification of *Krt19*-tdt^+^ cells in total AGER^+^ cells. In left image the quantification strategy is shown. Cells were identified based on co-localization of *Krt19*-tdt, AGER and DAPI (yellow doted lines) or AGER and DAPI (green dotted lines). Graph shows the percentage of KRT19^+^, AGER^+^ cells in total AGER^+^ cells in control and bleomycin injured mice on day-12. Data are from three independent experiments and are presented as mean ± s.e.m. Asterisk shows *p* < 0.0001. d, Immunostaining for CLU (green)/*Krt19*-tdTomato (red) in bleomycin-treated lungs. DAPI stains nuclei (blue). Scale bars indicate 30 µm.

**Supplementary Figure 7.**
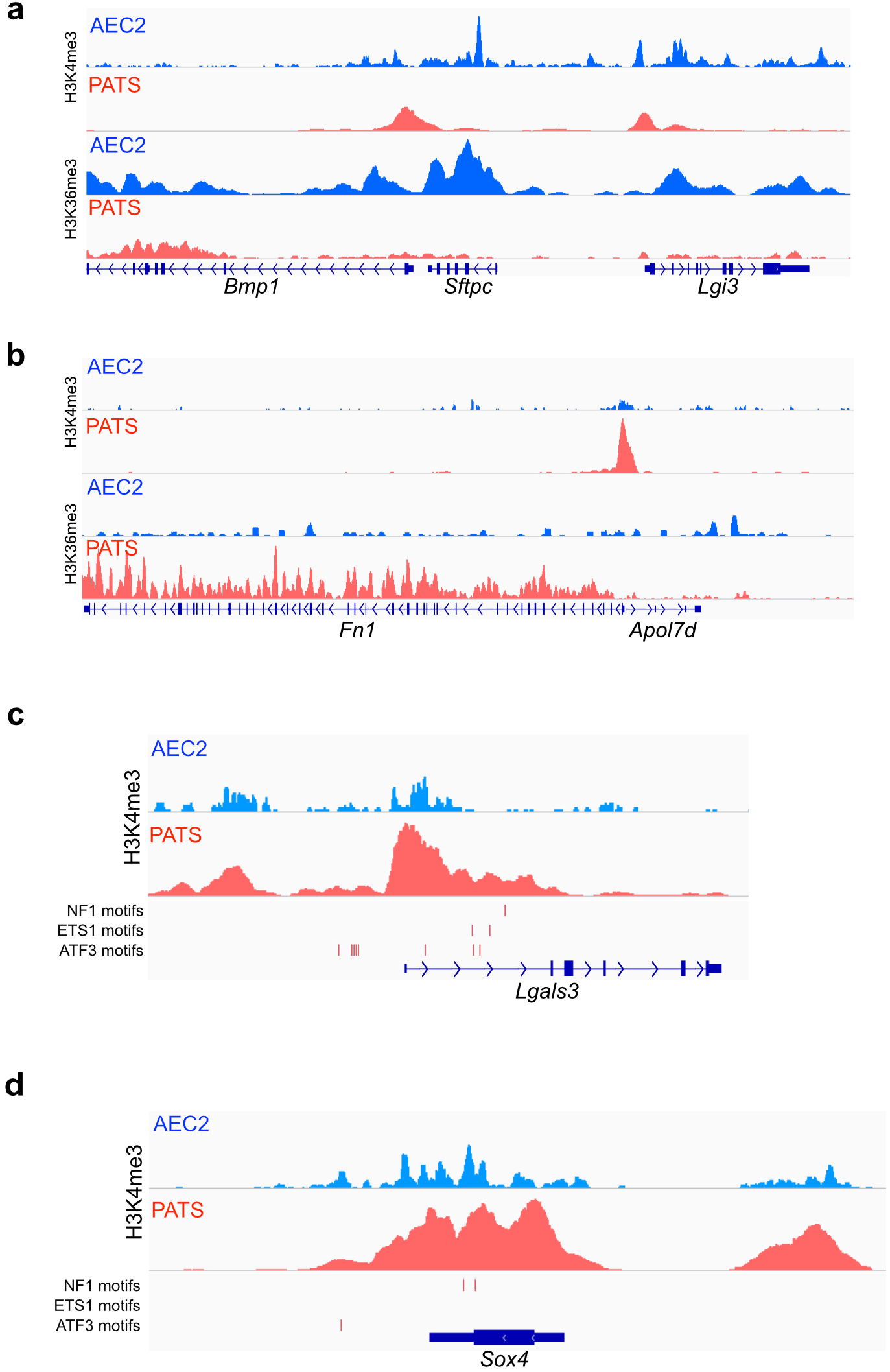
Histone marks profiling revealed active promoters and transcribing genomic loci in AEC2 and PATS following bleomycin-induced lung injury. Genomic tracks show enrichment of H3K4me3 and H3K36me3 in a), AEC2-specific gene loci (*Sftpc*) and b), PATS-specific gene loci (*Fn1*). H3K4me3 distribution and predicted binding motifs of NF1, ETS1 and ATF3 in c, *Lgals3* and d, *Sox4* loci. Blue and Red tracks indicate homeostatic AEC2 and PATS in bleomycin-induced lung injury, respectively. *y-* axis in AEC2 and PATS in all panels are normalized.

**Supplementary Figure 8.**
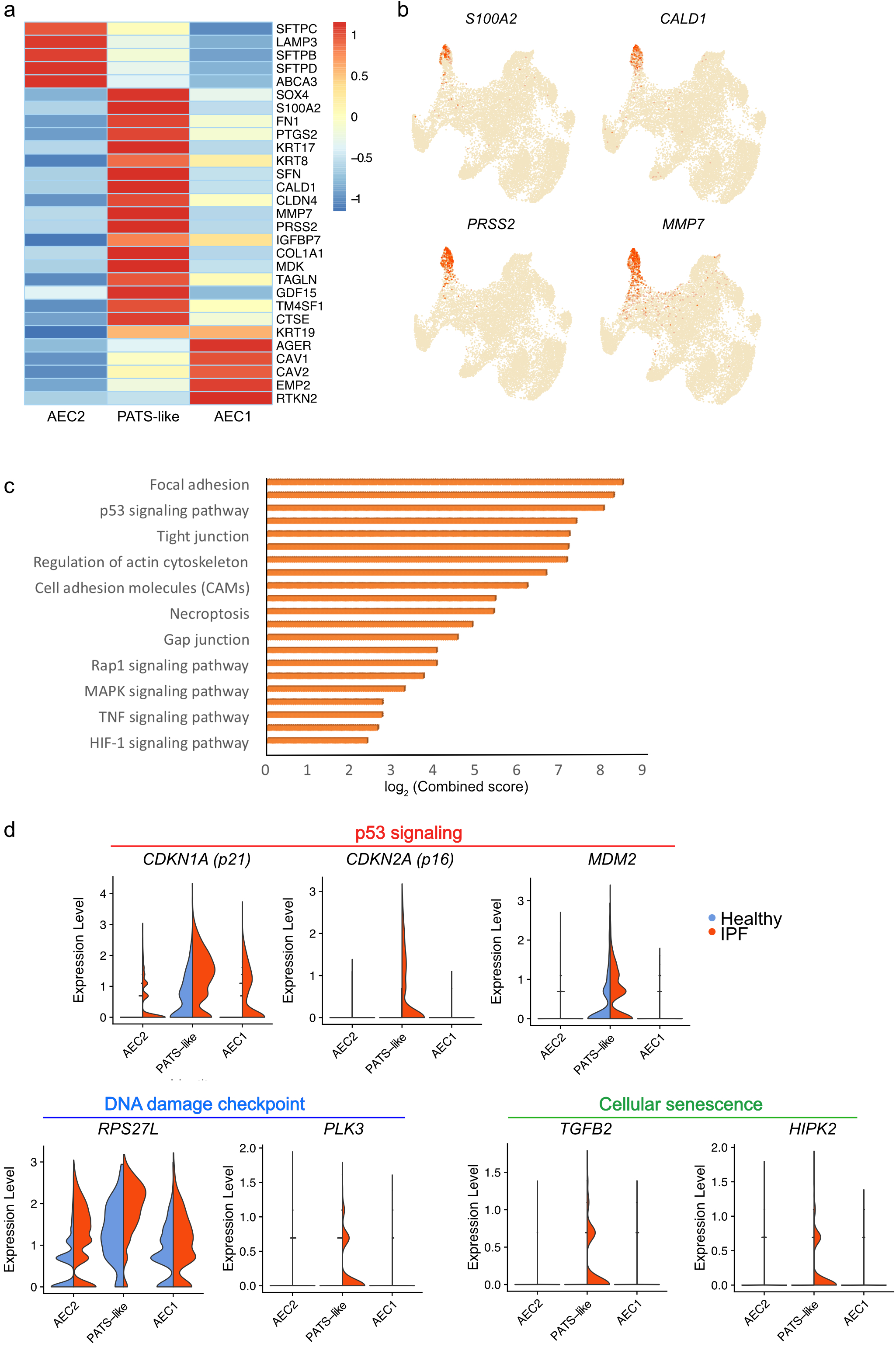
scRNA-seq analysis revealed the accumulation of PATS-like cells of human fibrotic lungs. a, Heatmap shows expression of marker genes of each cell population in human lungs (scale shows z-score). b, UMAP plots show the expression of indicated genes in alveolar epithelial populations in heathy controls and fibrotic human lungs. c, KEGG pathway enrichment analysis shows signaling pathways highly represented in PATS-like cells in human lungs. Scale shows log_2_ (combined score) obtained from Enrichr (see methods section for details). d, Violin plots shows the relative gene expression levels of indicated genes and cell types in control and IPF lungs.

**Supplementary Figure 9.**
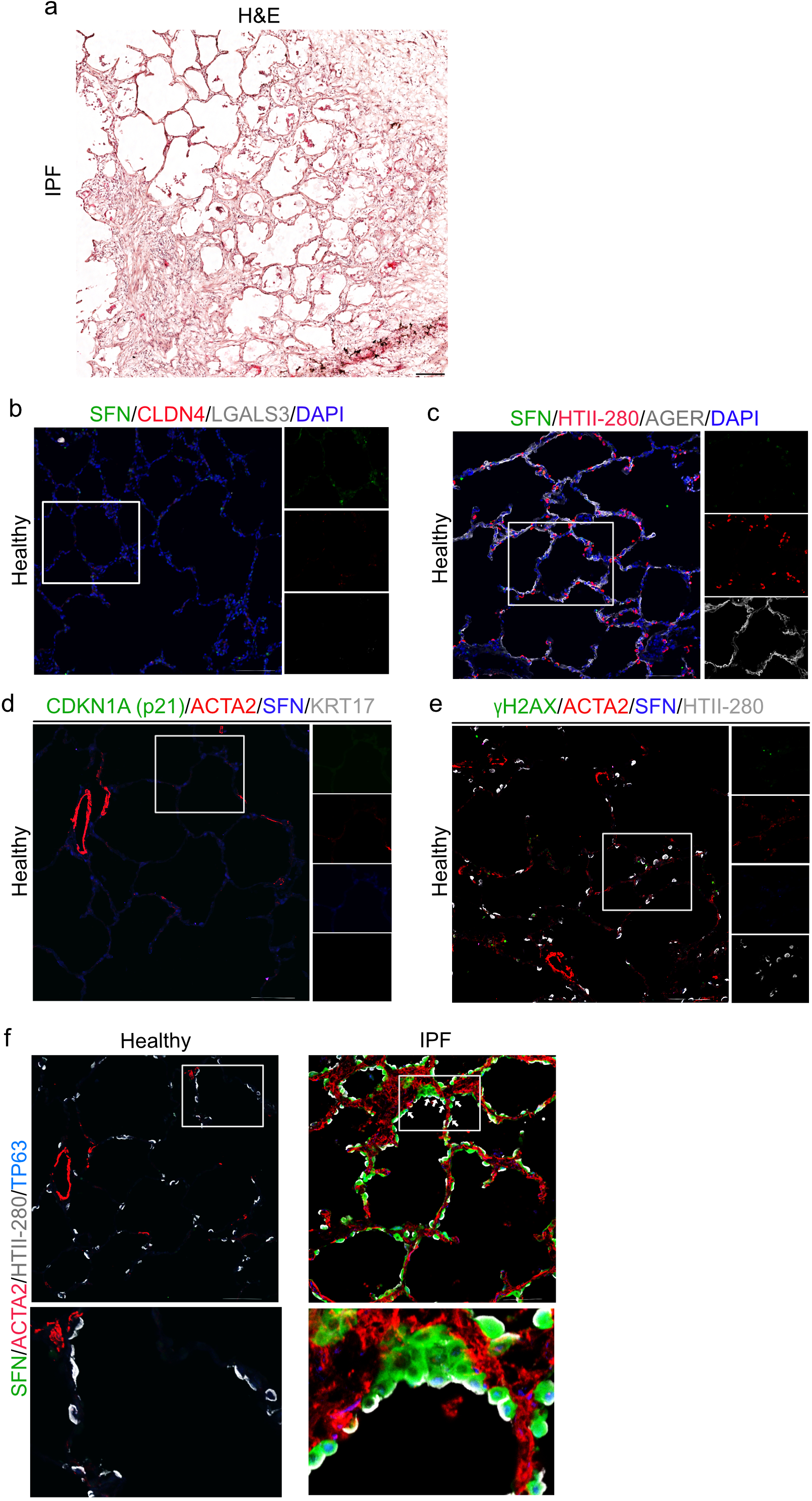
Markers specific to PATS-like cells are specifically expressed in human fibrotic lungs but not in healthy controls. a, Hematoxylin and Eosin staining of IPF lung section. Scale bar indicates 200 µm. b-e, Immunostaining for PATS-like markers in healthy human lungs. b, Co-staining for SFN (green), CLDN4 (red) and LGALS3 (grey), c, Immunostaining for SFN (green), HTII-280 (red) and AGER (grey), d, Co-staining for SFN (blue), p21 (green), ACTA2 (red) and KRT17 (grey) and e, Immunostaining for γH2AX (green), SFN (blue), ACTA2 (red) and HTII-280 (grey). White line box in merged images indicate region of single channel images shown on right. In f and g DAPI (blue) stains nuclei. f, Immunostaining for SFN (green), TP63 (blue), HTII-280 (grey) and ACTA2 (red) in healthy (left panel) and IPF lungs (right panel). Scale bars indicate 100 µm.

